# Inhibition mechanism of human sterol O-acyltransferase 1 by competitive inhibitor

**DOI:** 10.1101/2020.01.07.897124

**Authors:** Chengcheng Guan, Yange Niu, Si-Cong Chen, Yunlu Kang, Jing-Xiang Wu, Koji Nishi, Catherine C. Y. Chang, Ta-Yuan Chang, Tuoping Luo, Lei Chen

## Abstract

Sterol O-acyltransferase 1 (SOAT1) is an endoplasmic reticulum (ER) resident, multi-transmembrane enzyme that belongs to the membrane-bound O-acyltransferase (MBOAT) family ^1^. It catalyzes the esterification of cholesterol to generate cholesteryl esters for cholesterol storage ^2^. SOAT1 is a target to treat several human diseases ^3^. However, its structure and mechanism remain elusive since its discovery. Here, we report the structure of human SOAT1 (hSOAT1) determined by cryo-EM. hSOAT1 is a tetramer consisted of a dimer of dimer. The structure of hSOAT1 dimer at 3.5 Å resolution reveals that the small molecule inhibitor CI-976 binds inside the catalytic chamber and blocks the accessibility of the active site residues H460, N421 and W420. Our results pave the way for future mechanistic study and rational drug design of SOAT1 and other mammalian MBOAT family members.

## Main

Cholesterol is an essential lipid molecule in the cell membranes of all vertebrate. It is important for maintaining the fluidity and integrity of the membrane and is the precursor for the biosynthesis of other crucial endogenous molecules, such as steroid hormones and bile acids. In addition, cholesterol can modulate the activity of many membrane proteins such as GPCR ^4^ and ion channels ^5^. The concentration of cellular free cholesterol is highly regulated ^2^. Excessive intracellular cholesterol may form cholesteryl esters, which are catalyzed by the enzyme, sterol O-acyltransferase (SOAT), also called acyl-coenzyme A: cholesterol acyltransferase (ACAT). SOAT catalyzes the reaction between long chain fatty acyl-CoA and intracellular cholesterol to form the more hydrophobic cholesteryl ester, which is then stored in lipid droplets within the cell or transported in secreted lipoprotein particles to other tissues that need cholesterol. In addition to cholesterol, SOAT can use multiple sterols as substrates and activators ^3^. Because of its functional importance, SOAT1 is a potential drug target for Alzheimer’s disease ^6^, atherosclerosis ^7^ and several types of cancers ^8–11^.

Previous studies have shown that SOAT1 is an ER-localized multi-transmembrane protein that is evolutionary conserved from yeast to humans ^12^. There are two SOAT enzymes in mammals: SOAT1 and SOAT2, which have a protein sequence identity of 48% in human (258 out of 537 residues aligned). SOAT1 is ubiquitously expressed in many types of cells ^13^; while SOAT2 is mainly expressed in the small intestine and liver ^14^. Due to the pathophysiological importane of SOAT, many SOAT inhibitors of various strutural types have been made. Among them, the small molecule SOAT inhibitor CI-976 exhibit competitive inhibition against fatty acyl-CoA ^15^.

SOAT is the founding member of the membrane-bound O-acyltransferase (MBOAT) enzyme family, which transfers the acyl chain onto various substrates, including lipids, peptides and small proteins. There are 11 MBOAT family members in humans ^1^, which participate in many physiological processes, such as the last step of triglyceride biosynthesis catalyzed by acyl-CoA: diacylglycerol acyltransferase 1 (DGAT1), the maturation of hedgehog morphogen catalyzed by hedgehog acyltransferase (HHAT), and the acylation of peptide hormone ghrelin catalyzed by ghrelin O-acyltransferase (GOAT). Recently, the X-ray crystal structure of DltB from *Streptococcus thermophilus* was determined and is the only available structure of an MBOAT family member ^16^. The low-sequence conservation between DltB and SOAT (14.6% identity) hindered the accurate modeling of the human SOAT1 structure. Therefore, despite the important physiological functions of human SOAT enzymes, their architecture and mechanism remain elusive due to a lack of high-resolution structures. In this study, several human SOAT1 (hSAOT1) structures were determined. These structures reveal the architecture of hSOAT1, the binding sites of the competitive inhibitor CI-976, and provide a structural basis to understand the molecular mechanism of these enzymes.

### Structure determination

N-terminal GFP-tagged full-length (1–550) and N-terminal truncated hSOAT1 (66–550), solubilized in detergent micelles migrated before and after mouse TPC1 channel, which is a well-characterized dimeric channel with a molecular weight of 189 kDa ^17^ (Fig. S1a, S1b). This is consistent with previous studies, showing that hSOAT1 is a tetramer with a molecular weight of around 260 kDa ^18^ and that hSOAT1 became predominantly dimeric with the deletion of the N-terminal tetramerization domain ^19^. In order to measure the acyltransferase activity of purified protein, we developed the *in vitro* hSOAT1 assay using fluorescence-labeled NBD-cholesterol and oleoyl-CoA as substrates based on the previously reported SOAT1 whole cell assay ^20^ (Fig. 1a and Fig. S1c-h). The results show that the activity of hSOAT1 tetramer is linear within the first 15 min (Fig. S1h), and both the purified tetramer and dimer hSOAT1 protein exhibited cholesterol-activated O-acyltransferase activity in detergent micelles (Fig. 1b, Fig. S1d-h). Moreover, the SOAT enzyme activity is inhibited by CI-976 in a dose-dependent manner (Fig. 1c).

**Fig. 1.**
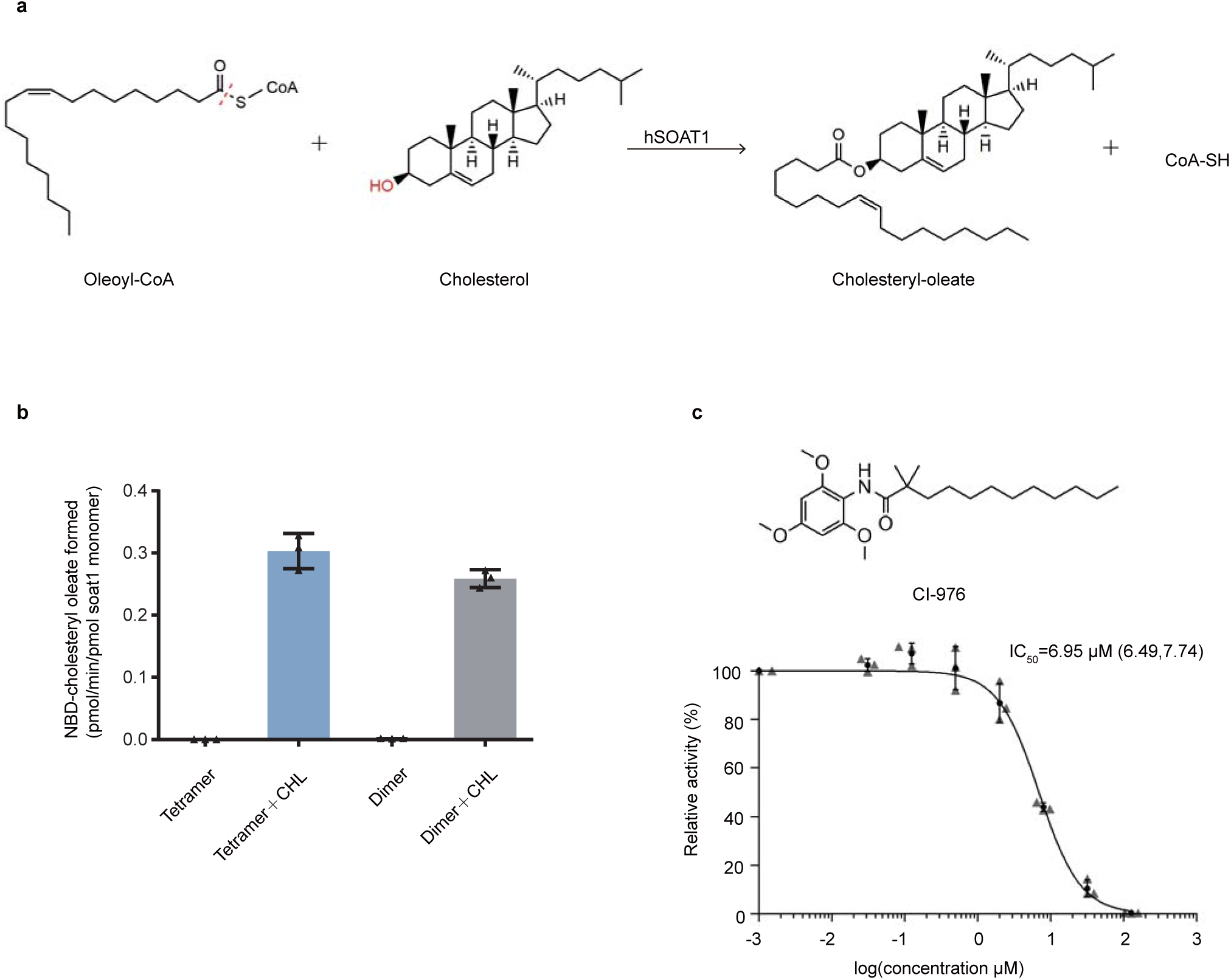
The enzymatic reaction catalyzed by hSOAT1. **a**, Chemical structures of the substrates and products of hSOAT1 enzyme are shown. The red dashed line indicates the bond that is broken during acyl-transfer reaction. The hydroxyl group that accepts acyl group is highlighted in red. **b**, The activation effect of cholesterol (CHL) on the esterification reaction of NBD-cholesterol catalyzed by hSOAT1 tetramer and dimer (Data are shown as means ± standard deviations, n = 3 biologically independent samples). **c**, Chemical structure of CI-976 and dose-dependent inhibition curve of hSOAT1 tetramer by CI-976 (The first data point is an artificial point. Data are shown as means ± standard deviations, n = 3 biologically independent samples, and numbers in parentheses are the range for IC50 obtained from curve fitting).

To investigate the structure of hSOAT1, we prepared the sample of hSOAT1 in the presence of CI-976 in detergent micelles for cryo-EM analysis. The 3D classification showed the hSOAT1 protein exhibited severe conformational heterogeneity (Fig. S2). One of the 3D classes can be further refined to 12 Å and the map allowed the visualization of the general shape of hSOAT1, in which the cytosolic N terminal oligomerization domain lays above the transmembrane domain. Moreover, the central slice of the transmembrane domain density map indicated the hSOAT1 is a tetramer composed of dimer of dimers (Fig. S2d), which is consistent with previous biochemical data ^19^. To further stabilize the transmembrane domain, we reconstituted hSOAT1 into a lipid nanodisc (Fig. S3a, S3b) for cryo-EM analysis (Fig. S4). The top view of 2D class averages of the nanodisc sample had markedly enhanced features and confirmed the transmembrane region of hSOAT1 to be a dimer of dimers (Fig. S4b). However, some 2D class averages in top views showed that one distinct dimer was adjacent to another blurry but still distinguishable dimer (Fig. S4b), suggesting the dimer-dimer interface is mobile. Through multiple rounds of 2D and 3D classification, two 3D classes with discernable transmembrane helix densities were isolated. Subsequent refinements generated reconstructions at resolutions of 8.2 Å (for the oval-shaped tetramer) and 7.6 Å (for the rhombic-shaped tetramer) (Fig. S4c, S4d). The oval-shaped structure occupies a 3D space of 170 Å×120 Å×50 Å and the shape is similar to 3D Class 1 observed in detergent micelles (Fig. S2), and the rhombic-shaped structure occupies 185 Å×110 Å×50 Å (Fig. 2), similar to the 3D Class 2 observed in detergent micelles (Fig. S2). The N terminal tetramerization domain is invisible in both maps probably due to its flexibility. These two structures reveal that the dimer-dimer interfaces of oval-shaped and rhombic-shaped hSOAT1 are distinct (Fig. 2b, d), correlating with the 3D heterogeneity observed in detergent micelles. This suggests the interfaces between dimers are unstable and dynamic in nature, which in turn hampered high-resolution structure determination. To overcome the conformational heterogeneity in the tetrameric hSOAT1 sample, the functional dimer construct (hSOAT1 66-550) was expressed and purified. The protein was reconstituted into the nanodisc in the presence of inhibitor CI-976 (Fig. S3c, S3d). Subsequent 3D reconstruction generated a 3.5 Å cryo-EM map, which allowed the model to be built *de novo* (Fig. 3a, Fig. S5-6, and Table S1).

**Fig. 2.**
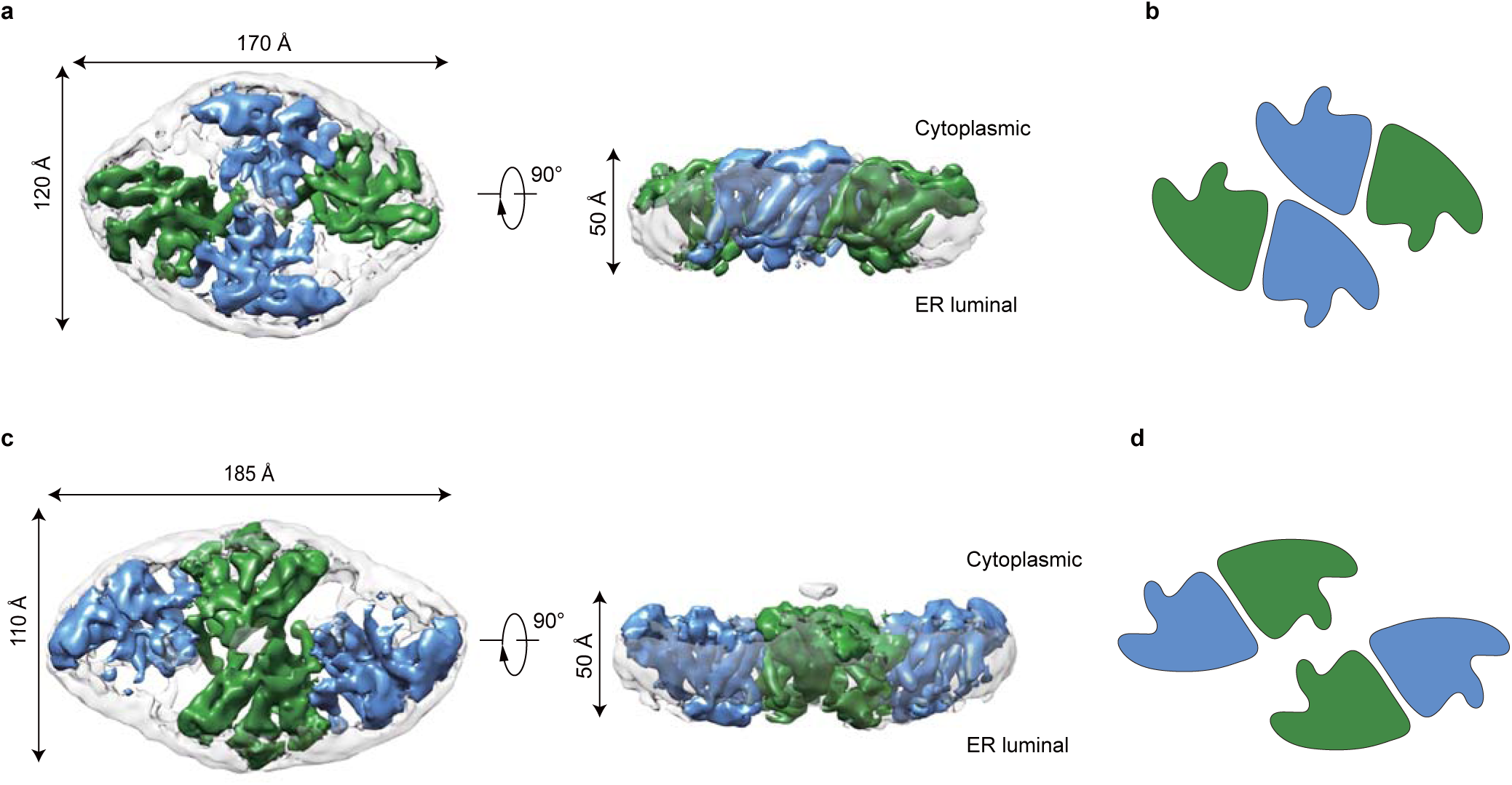
Cryo-EM maps of the human SOAT1 tetramer. **a**, Cryo-EM density map of the oval-shaped hSOAT1 tetramer in top view and side view. Two subunits in one hSOAT1 dimer are colored in green and blue, respectively. Densities of the MSP and lipids in nanodiscs are colored in gray with semi-transparency. **b**, The domain arrangement of oval-shaped hSOAT1 tetramer is shown in a cartoon model in top view. Each subunit is colored the same as in (**a**). **c**, The density map of the rhombic-shaped hSOAT1 tetramer is shown in top and side view. **d**, The domain arrangement of rhombic-shaped hSOAT1 tetramer in top view. Each subunit is colored in the same way as in (**c**).

**Fig. 3.**
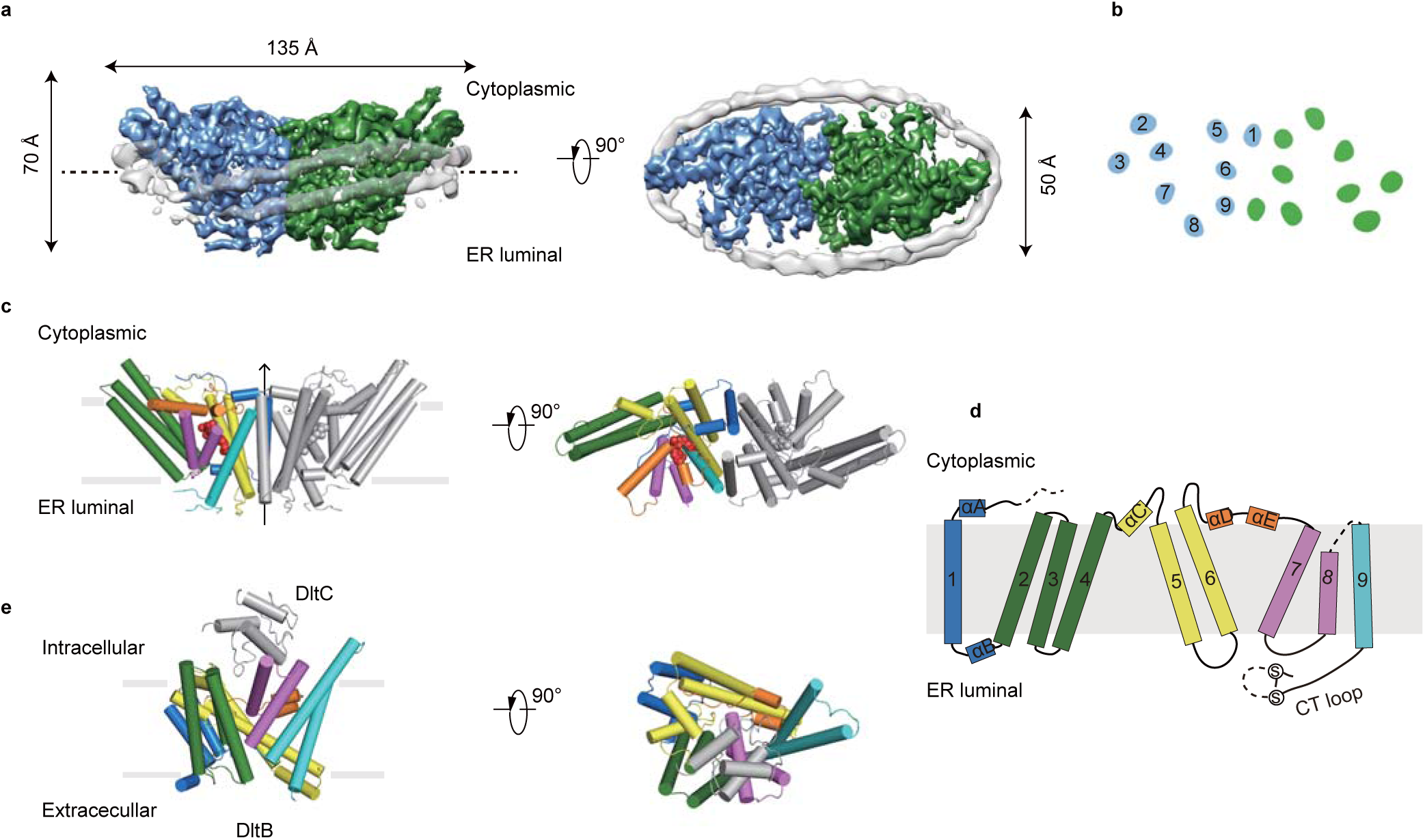
The structure of hSOAT1 dimer. **a**, Cryo-EM map of hSOAT1 dimer in side view and top view. Two subunits of the dimer are colored in green and blue. Density corresponding to the nanodisc is colored in gray with semi-transparency. **b**, Top view of the cross-section of the transmembrane domain at the approximate level indicated by the dashed lines in (**a**). The identities of the transmembrane helices from the blue subunit are labeled in numbers. **c**, The structural models of hSOAT1 dimer are shown in side view and top view. Helices are shown as cylinders. One subunit of hSOAT1 is in rainbow color and the other subunit is in grey. CI-976 molecule is shown as red spheres. **d**, The topology of one hSOAT1 subunit. The colors are used in the same way as in (**c**). **e**, The crystal structure of DltB-DltC complex in side view and top view (PDB ID: 6BUG). The DltB subunit is in rainbow color while the DltC subunit is in gray.

### Structure of hSOAT1 dimer

The hSOAT1 dimer has a symmetric “rubber raft” shape and occupies a 135 Å×70 Å×50 Å three-dimensional space (Fig. 3a). Each hSOAT1 subunit is composed of nine transmembrane helices (Fig. 3b), which is consistent with previous prediction based on the biochemical data ^21^. The first 52 amino acids (66-117) of the dimeric hSOAT1 were invisible in the cryo-EM map, presumably due to their high flexibility (Fig. 3c, d). Notably, this region has the low sequence conservation between SOATs from different species and paralogues. The cytosolic pre-αA loop runs parallel to the membrane. The amphipathic αA floats on the putative lipid bilayer with hydrophobic di-leucine motifs facing the membrane and connects to M1 via a near 90° turn (Fig. 3d). M1 is linked to the long tilted M2-M4 tri-helix bundle by an ER-luminal loop and a short helix αB (Fig. 3d). The six transmembrane helices M4-9 form a funnel-shaped central cavity, which is capped by the cytosolic helices αC, αD, and αE on the top. At the end of M9, a C-terminal ER luminal loop (CT loop) is cross-linked by the disulfide bond formed between C528 and C546 (Fig. 3d). It was previously reported that this loop is important for hSOAT activity and stability ^22,23^. Indeed, the CT loop interacts with both the luminal M7-M8 loop and the M5-M6 loop (Fig. 3d) at the ER luminal side. Based on the sequence homology (Fig. S7), the closely related SOAT2 and the distal related DGAT1 enzymes might have a similar transmembrane domain topology to hSOAT1 reported here. In addition, the TM2-9 of hSOAT1 share a similar structural fold with H4-H16 of DltB ^16^, with core RMSD of 3.2 Å for 227 structurally aligned residues, despite of their low sequence identity (Fig. 3e), suggesting a common evolutionary origin of the MBOAT family enzymes.

### Dimer interface

In contrast to the monomeric DltB protein, the functional unit of hSOAT1 is a tightly packed dimer, with a dimer interface area of 6,520 Å^2^. The two hSOAT1 protomers interact through the M1, M6, M6-αD loop and M9 helices in a symmetric way (Fig. 4a and b). In the inner leaflet of the ER membrane, M144, A147, L148 and L151 on M1 of one subunit interact with I370 and F378 on M6, and V501, W504 and F508 of M9 of the other subunit (Fig. 4c and d). In the outer leaflet of ER membrane, V158 and V159 on M1 interact with V363 on M6 of the other subunit (Fig. 4c and d). The residues that form the dimer interface are mostly hydrophobic and interact with each other in a shape-complementary manner.

**Fig. 4.**
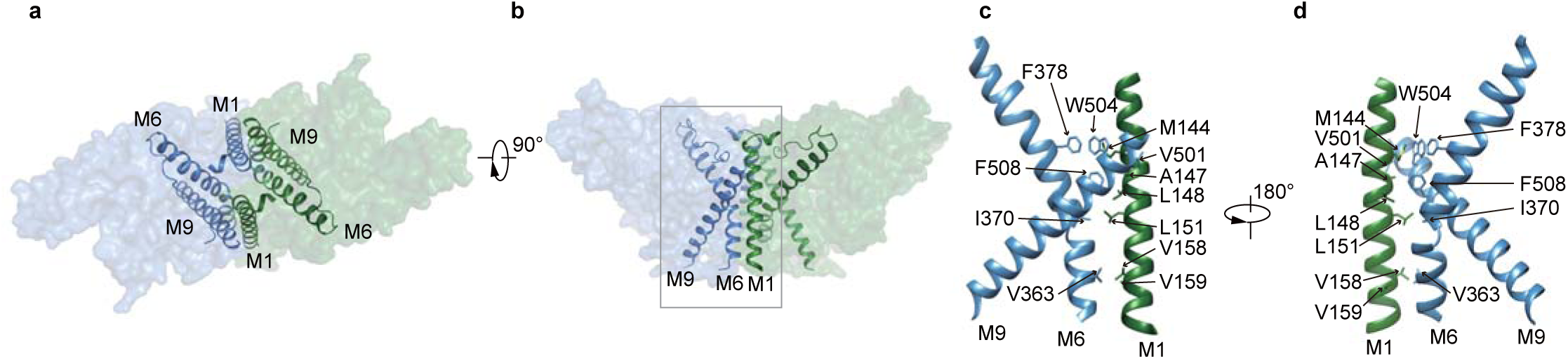
The dimer interface of hSOAT1 dimer. **a-b**, Top view and side view of one hSOAT1 dimer are shown in surface representation with semi-transparency. M1, M6 and M9 helices are shown as ribbons. **c**, Close-up view of the dimer interface boxed in (**b**), with interacting residues shown in sticks. **d**, A l80° rotated view compared to (**c**).

### Reaction chamber of hSOAT1

Previous studies suggest that the conserved H460 on M7 is crucial for hSOAT1 activity and is the putative catalytic residue ^21^. The side chain of H460 points towards the interior of the central cavity cradled by M4−M9 (Fig. 5a and b). These structural observations suggest this central cavity is the chamber where the acyl transfer reaction takes place. In accordance with this, previous studies have also identified several residues in the central cavity that are important for hSOAT1 function. Mutations of residues on M7 and M8 affect the catalytic activity ^24,25^. C467 at the end of M7 is the major target site for p-chloromercuribenzene sulfonic acid-mediated SOAT1 inactivation ^26^. These data also indicate that the local environment in the central cavity is important for the catalytic reaction.

**Fig. 5.**
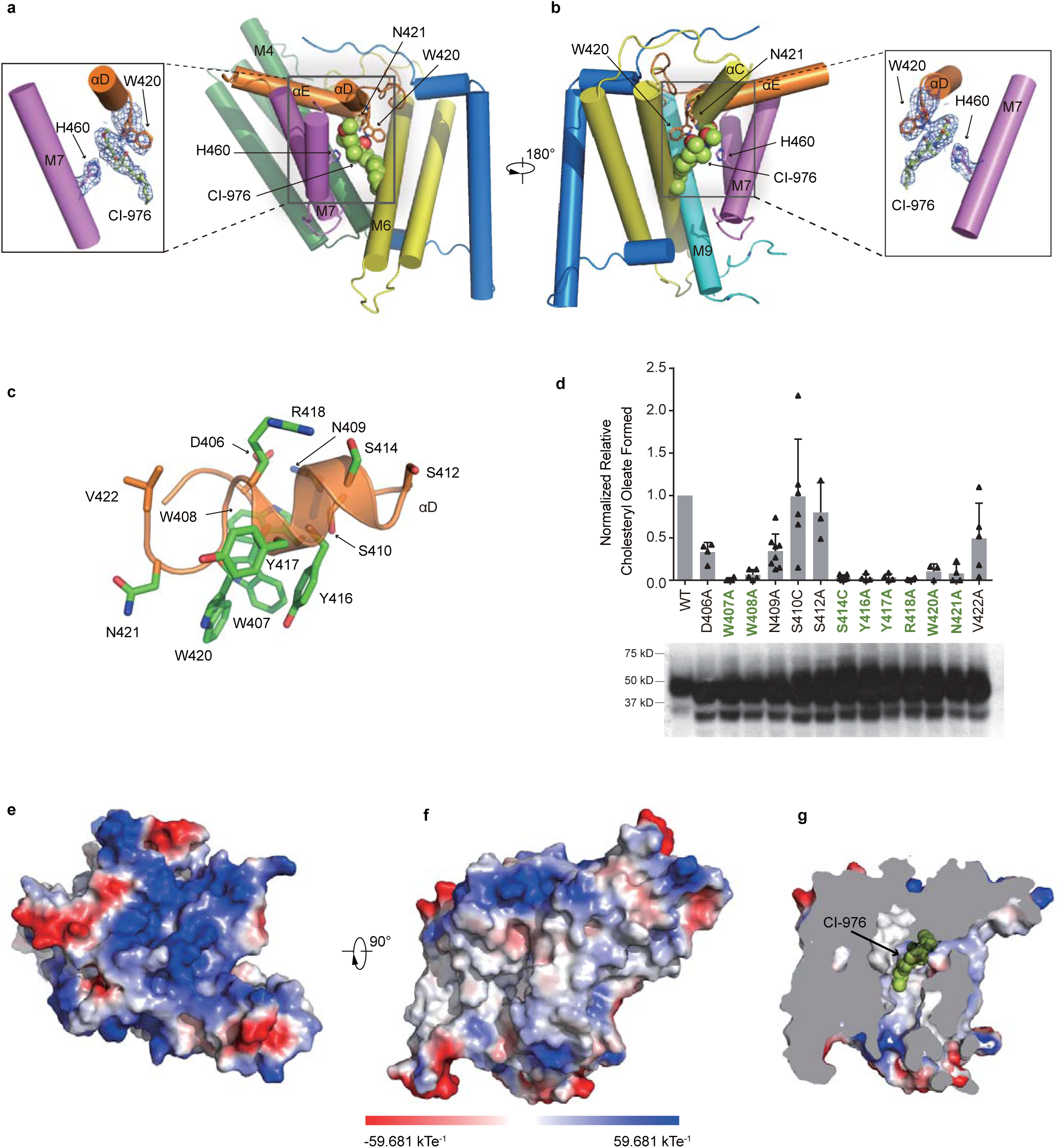
The catalytic chamber and the CI-976 binding site in hSOAT1. **a-b**, Side views of one hSOAT1 subunit in cartoon representation. The transmembrane helices are colored in the same way as in Fig. 3c, and **3d**. The inhibitor CI-976 is shown in lemon sphere. The side chains of residues that are close to CI-976 are shown in sticks. In the inlet, the cryo-EM densities of CI-976 and adjacent side chains were shown in blue meshes at the same contour level. Maps were further sharpened at −50 Å by Coot for visualization. **c**, Close-up view of the M6-αD loop and αD with side chains shown in sticks. Residues that are important for SOAT1 enzyme activity reported in (**d**) were colored in green. **d**, Enzymatic activities of various hSOAT1 mutants. (For D406A, n=4. For W407A, n=4. For W408A, n=5. For N409A, n=8. For S410C, n=6. For S412A, n=3. For S414C, n=6. For Y416A, n=4. For Y417A, n=4. For R418A, n=4. For W420A, n=3. For N421A, n=4. For V422A, n=5.) Data are shown as means ± standard errors. A two-tailed unpaired t test was used to calculate the p values. For S410C, p=0.9434. For S412A, p=0.017. Other mutants had p values less than 0.0001. The bottom showed the western results of the hSOAT1 mutants proteins. **e-f**, The top and side views of one hSOAT1 monomer in the surface representation. The surfaces are colored by electrostatic potential calculated by Pymol. **g**, The cut-away view showing the binding pocket of the inhibitor CI-976 inside the reaction chamber.

The reaction chamber is covered by a lid formed by M4-αC loop, αC, M6-αD loop, αD and αE on the cytosolic side. It is suggested that part of the cytosolic lid, residues 403-409 on M6-αD loop, may be involved in the binding of fatty acyl-CoA ^27^. To further explore the role of the residues in this region, we performed alanine or cysteine mutagenesis of conserved residues within region amino acids 406-422 (from M6-αD loop to αD, Fig. 5c), and analyzed the effects of mutations after transient expressions of each of these mutant plasmid DNAs in a CHO cell clone AC29, that is devoid of endogenous SOAT activity, but regains enzyme activity upon transient expression of hSOAT plasmid DNA ^28^. The results showed that mutating any of the following residues W407, S414, Y416, Y417, R418, W420, and N421 to alanines or cysteines caused loss in normalized hSOAT1 enzyme activity (by greater than 90%) without severely lowering the cellular hSOAT1 protein expression in transfected cells (Fig. 5d). Because the hydrophilic side chains of S414 and R418 on αD face the cytosolic side of SOAT1, S414 and R418 might be involved in functions important for hSOAT1 activity, such as binding of highly hydrophilic CoA group of fatty acyl-CoA substrate. These results further emphasize the important role of the cytosolic lid of the reaction chamber in the enzymatic reaction.

### Inhibitor and ligands binding sites

A strong extra non-protein density was found in the central cavity and the size and shape of the density matched that of the inhibitor CI-976, which was included during cryo-EM sample preparation (Fig. S8a). By comparing the current map with the maps without CI-976 (as described later), we proposed that this “nonprotein material” represents the CI-976 molecule. The large trimethoxyphenyl group of CI-976 is sandwiched between the catalytic residue H460 on M7, and residues N421 as well as W420 on the αD-αE loop (Fig. 5a, b), all of which are crucial for the catalytic activity of hSOAT1. The elongated dodecanamide tail of CI-976 extends in the cavity and interacts with M6 and M9. The binding position of CI-976, right in the catalytic center, suggests that it inhibits the enzyme by precluding substrate loading into the catalytic center, which is consistent with the competitive behavior of CI-976 ^15^. Moreover, it has been reported that certain residues on M9 are responsible for the selectivity of subtype-specific SOAT inhibitors, such as pyripyropene A ^29^. This further suggests that the catalytic chamber might be a common binding site for inhibitors with diverse chemical structures.

There are several extra non-protein densities observed in the cryo-EM map. One density (density A) is close to the dimer interface. It is surrounded by L129, L132 and L133 on the hydrophobic side of αA, F145 on M1, C333 on M5, F382 on M6 and W408 on M6-αD loop (Fig. S8c). The shape of this density is close to cholesterol, suggesting this ligand might be a tightly-bound sterol-like molecule that was carried on during membrane protein extraction and purification. SOATs are allosteric enzymes that can be activated by cholesterol ^3^ and it is predicted that SOATs have two functional distinct cholesterol binding sites. One site is the substrate binding site and the other is the allosteric activating site that provides the feedback regulation mechanism regarding cholesterol concentration in the ER ^3^. On the other hand, previous work showed that SOAT1 exhibits only low affinity binding towards cholesterol, either as substrate or as activator, with dissociation constant at sub-millimolar concentration ^30^. Therefore, we speculate that the molecule in density A is neither a cholesterol substrate nor a cholesterol activator, but a sterol-like molecule bound to the enzyme in the sample preparation procedure. Another elongated density (density B) is inside the central cavity and surrounded by F258 and R262 on M4, F384 and W388 on M6, P304 on αC-M5 linker and V424 on αE (Fig. S8e). One additional density (density C) is on the ER luminal side of the central cavity and surrounded by Y176 on αB, S519, W522 and Y523 on M9, L468 on M7-M8 linker, P250 on M3−M4 linker (Fig. S8g). The exact identities of these densities A-C and their roles on the hSOAT1 function remain elusive.

In order to gain more mechanistic insights into the catalytic mechanism of hSOAT1 and to trap the catalytic reaction in a transition state, we designed and synthesized a compound that might mimic the catalytic transition intermediate. Inspired by the previous work on GOAT ^31^, we hypothesized that the catalytic reaction intermediate of hSOAT1 might be a ternary complex of sterol, acyl-CoA and the enzyme. Therefore, the covalent linkage of sterol and acyl-CoA would yield a competitive inhibitor with a higher affinity than each individual substrate alone. Pregnenolone was previously reported to be a substrate of hSOAT1 with a lower K_m_ and better solubility than cholesterol ^32^. In the current study, CoA group was chemically covalently linked with stearoyl-pregnenolone to generate a bi-substrate analogue for hSOAT1 (BiSAS) (Fig. S9a). Indeed, BiSAS inhibited the purified hSOAT1 enzyme in the *in vitro* NBD-cholesterol based assay (Fig. S9b). A cryo-EM sample of hSOAT1 dimer was prepared in the presence of BiSAS and cholesterol (Fig. S3e, S3f) and the cryo-EM reconstruction generated a 3.5 Å map (Fig. S10-11 and Table S1). This map was compared with the CI-976 bound map and was found to be similar overall, with a real space correlation of 0.9. We anticipated that BiSAS might mimic both substrates of hSOAT1 and might occupy the substrate-binding pocket while cholesterol might only bind at the activating site. In contrast to our prediction, in the BiSAS map, the strong continuous density of CI-976 found in the central cavity of the CI-976 bound map was replaced by weak residual densities that were not continuous (Fig. S8b), indicating the absence of full-sized BiSAS molecule, probably due to the low affinity or incompatibility of BiSAS in the nanodisc sample preparation conditions. Retrospectively, the large size of BiSAS molecule would not fit into the hSOAT1 structure in the current conformation. In addition, we did not observe any extra density that would suggest the presence of activating cholesterol either, probably due to the low affinity of activating cholesterol on hSOAT1 ^30^. Instead, most likely, the BiSAS map represents the apo resting state of hSOAT1. Interestingly, all three additional non-protein ligand densities (density A−C) present in the CI-976 map were also observed in the BiSAS map (Fig. S8d, S8f, S8h), further suggesting their tight associations with the hSOAT1 protein and likely their functional importance as well.

### A working model for hSOAT1 activation

SOAT catalyzes the esterification reaction between acyl-CoA and cholesterol. The surface representation of the hSOAT1 monomer shows an intra-membrane tunnel from outside of the molecule into the reaction chamber. The tunnel is located between M4 and M5, and is mainly hydrophobic. This lateral tunnel within the transmembrane domain of hSOAT1 might be the substrate or product transfer pathway for hydrophobic molecules. The other substrate, fatty acyl-CoA, is amphipathic with a hydrophobic tail and a highly hydrophilic CoA group. The cytosolic acyl-CoA can access the central reaction chamber only from the cytosolic side of hSOAT1. However, the surface representation of the hSOAT1 dimer shows that the reaction chamber is completely shielded from the cytosolic side by the two short αD-αE helices and associated intracellular loops (Fig. 5e-g). This suggests that the current structure represents a resting state with relatively low catalytic activity, in which the putative catalytic residue H460 is less accessible to the acyl-CoA substrate (Fig. S11). Therefore an activation step that opens the reaction chamber, probably caused by sterol binding at the allosteric activator site, is required for the sterol-dependent fully activation of hSOAT1 ^4^.

The structures of human SOAT1 presented here provide the a high-resolution view of the architecture and domain organization of this important enzyme and shed light on the structure of other closely related MBOAT family proteins, such as SOAT2 and DGAT1. This work not only paves the way towards a better mechanistic understanding of SOAT1-catalyzed reaction, but also provides a template for structure-based inhibitor design to target several human diseases.

## Acknowledgement

The cDNAs of human SOAT1 and SOAT2 were kindly provided by Jiahuai Han. Cryo-EM data collection was supported by Electron microscopy laboratory and Cryo-EM platform of Peking University with the assistance of Xuemei Li, Daqi Yu, Xia Pei, Bo Shao, Guopeng Wang, and Zhenxi Guo. Part of structural computation was also performed on the Computing Platform of the Center for Life Science and High-performance Computing Platform of Peking University. This work is supported by grants from the Ministry of Science and Technology of China (National Key R&D Program of China, 2016YFA0502004 to L.C.), National Natural Science Foundation of China (91957201, 31622021, 31870833 and 31821091 to L.C., 31521004, 21672011 and 21822101 to T. L.), Beijing Natural Science Foundation (5192009 to L.C.), and Young Thousand Talents Program of China to L.C, NIH grants in U.S.A. (R01AG037609 and AG063544 to T.Y. Chang and C.C.Y. Chang). C.G. is supported by a Boehringer-Ingelheim postdoctoral fellowship.

## Author Contribution

Lei Chen initiated the project. Chengcheng Guan developed the fluorescence-based activity assay. Chengcheng Guan purified proteins and prepared the cryo-EM samples with the help of Yange Niu. Chengcheng Guan collected the cryo-EM data with the help of Jing-Xiang Wu and Yunlu Kang. Chengcheng Guan processed the cryo-EM data with the help of Lei Chen. Lei Chen built and refined the atomic model. Si-Cong Chen synthesized the BiSAS inhibitor under the guidance of Tuoping Luo. Koji Nishi, Catherine C. Y. Chang and Ta-Yuan Chang did the enzymatic activity assays based on radioactive substrates. All authors contributed to the manuscript preparation.

## Materials & Correspondence

Correspondence to Lei Chen. Any materials that are not commercially available can be obtained upon reasonable request.

## Data availability

The cryo-EM map of hSOAT1 tetramer in oval shape, hSOAT1 tetramer in rhombic shape, hSOAT1 dimer bound with CI-976 and in apo resting state have been deposited in the EMDB under ID codes EMD-0829, EMD-0830, EMD-0831 and EMD-0832. The atomic coordinates of hSOAT1 dimer bound with CI-976 and in apo resting state have been deposited in the PDB under ID codes 6L47 and 6L48.

## Methods

### Cell culture

Sf9 insect cells were cultured in Sf-900 III serum-free medium (SFM; Thermo Fisher Scientific) or in SIM SF (Sino Biological) at 27 °C. HEK293F cells were cultured at 37 °C with 6% CO2 and 70% humidity in Free Style 293 medium (Thermo Fisher Scientific) supplemented with 1% fetal bovine serum (FBS).

### Chemical synthesis of BiSAS

To the solution of α-bromo stearic acid (727 mg, 2 mmol, 1.0 equiv), pregnenolone (632 mg, 2 mmol, 1.0 equiv) and dicyclohexylcarbodiimide (DCC, 495 mg, 2.4 mmol, 1.2 equiv) in dichloromethane (DCM, 30 mL) was added 4-dimethylaminopyridine (DMAP, 293 mg, 2.4 mmol, 1.2 equiv). The solution was stirred at room temperature for 24 h. The crude product was purified by column chromatography (Hexanes: Ethyl acetate = 10:1) to obtain the α-bromo ester (747 mg, 57 % yield) as a white solid. ^1^H NMR (400 MHz, CDCl_3_) *δ* 5.39 (d, J = 5.1 Hz, 1H), 4.67 (qd, J = 11.3, 9.5, 4.2 Hz, 1H), 4.17 (t, J = 7.4 Hz, 1H), 2.54 (t, J = 8.9 Hz, 1H), 2.40 – 2.32 (m, 2H), 2.12 (s, 4H), 2.08 – 1.84 (m, 6H), 1.75 – 1.58 (m, 4H), 1.56 – 1.39 (m, H), 1.37 – 1.11 (m, 32H), 1.03 (s, 3H), 0.88 (t, J = 6.7 Hz, 3H), 0.63 (s, 3H). ^13^C NMR (101 MHz, CDCl_3_) *δ* 209.69, 169.49, 139.39, 122.83, 77.48, 77.16, 76.84, 75.51, 63.81, 56.96, 49.99, 46.72, 44.12, 38.92, 37.69, 37.04, 36.74, 35.04, 32.08, 31.94, 31.91, 31.71, 29.84, 29.83, 29.81, 29.73, 29.61, 29.52, 29.45, 28.97, 27.61, 27.40, 24.62, 22.97, 22.84, 21.18, 19.46, 14.28, 13.37. HRMS(ESI): m/z calcd for C_39_H_66_BrO_3_^+^ [M + H]^+^: 661.418985, found 661.420600

To the solution of α-bromo ester obtained above (27 mg, 40 μmol, 1.0 equiv.) and CoA-SH (62 mg, 80 μmol, 2.0 equiv) in *N,N*-dimethyllformamide (DMF, 1 mL) was added triethylamine (TEA, 56 μL, 0.4 mmol, 10 equiv). The solution was stirred under nitrogen atmosphere at 35 °C overnight. The crude product was purified by reverse phase HPLC (Water : Acetonitrile = 50 : 50 to 5 : 95) to obtain BiSAS in bis(triethylammonium) salt form (23.8 mg) as a colorless solid. ^1^H NMR (500 MHz, D_2_O) *δ* 8.44 (s, 1H), 8.06 (s, 1H), 6.04 (s, 1H), 5.27 (s, 1H), 4.89 – 3.72 (m, 9H), 3.65 – 3.20 (m, 6H), 3.10 (q, J = 7.2 Hz, 8H), 2.89 – 1.73 (m, 19H), 1.42 – 1.00 (m, 48H), 0.88 (s, 3H), 0.77 (s, 6H), 0.72 – 0.60 (m, 3H), 0.50 (s, 3H). HRMS(ESI): m/z calcd for C_60_H_99_N_7_O_19_P_3_S^-^ [M - H]^-^: 1346.593480, found 1346.591330.

### Constructs

We cloned human SOAT1 (Hs_SOAT1), human SOAT2 (Hs_SOAT2), Xenopus laevis SOAT1 (Xl_SAOT1), chicken SOAT1 (Gg_SAOT1) and zebrafish SOAT1 (Dr_SOAT1) into NGFP tagged BacMaM vector for screening by FSEC methods ^33^. The screening procedures identified human SOAT1 as a putative target with reasonable expression level and elution profile. cDNAs of human SOAT1 full-length (1-550), SOAT1-dimer (66-550) were cloned into a modified BacMaM vector, with N-terminal His_7_-strep-GFP tags ^34^.

### Protein expression and purification

The BacMam expression system was used for large-scale expression of human SOAT1. The BacMaM viruses were added into suspended HEK293F cells (grown in FreeStyle 293 medium + 1% FBS, 37°C). Sodium butyrate (10 mM) was added to the culture 12 hours post-infection to promote protein expression and the temperature was lowered to 30°C. Cells were harvested 60 hours post-infection and washed with TBS buffer (20 mM Tris pH 8.0 at 4°C, 150 mM NaCl). Cell pellets were frozen at −80°C for later use.

The cell pellets were resuspended in TBS buffer supplemented with protease inhibitors (2µg/ml leupepetin, 2µg /ml pepstatin, 2µg /ml aprotonin and 1 mM PMSF). Unless stated otherwise, all buffers used for purification were supplemented with inhibitor either 1μM inhibitors CI-976 or BiSAS. The cells were broken by sonication and centrifuged at 8,000 rpm for 10 min with JA25.5 rotor (Beckman) to remove cell debris. The supernatant was centrifuged at 40,000 rpm for 1 h in Ti45 rotor (Beckman) to harvest cell membrane in pellets. The membrane pellets were homogenized in TBS, solubilized by 1% digitonin for 2 h at 4°C, and centrifuged at 40,000 rpm for 1 h. The supernatant was loaded onto a 1 ml prepacked strep-tactin superflow high-capacity column (IBA) and washed using TBS buffer with 0.1% digitonin. The binding protein was eluted using TBS buffer with 0.1% digitonin and 10 mM desthiobiotin. The eluted proteins were digested with PreScission protease to remove tags and further purified by superose-6 increase column (GE Healthcare) in TBS buffer with 0.1% digitonin or 40 μM GDN. Buffers were not supplemented with inhibitors during the purification of proteins used for the activity assay. Purified protein were either directly used for experiments or flash frozen in liquid nitrogen for storage at −80°C. Stored protein was thawed on ice and centrifuged to remove precipitates before further experiments.

### SOAT1 activity assays using ^3^H-oleate in intact cells

This assay was designed to measure the rate of ^3^H-cholesteryl oleate biosynthesis in intact cells. Mutant CHO AC29 cells that lack endogenous SOAT activity were cultured in 6-well plates at 37 °C and transiently transfected with DNAs from various single amino acid substitutions of hSOAT1 as indicated. Unless stated otherwise, each construct contained the 6x His tag at the N-terminus. We used hSOAT1 that contained the C92A substitution as the “wild type” hSOAT1. Control experiments showed that the SOAT activity of the C92A mutant remained the same as the wild type ^21^ but the C92A substitution greatly diminished the SOAT1 protein aggregation that occurred *in vitro* during the SDS-PAGE process.

At the third day after transfections, the cells were split into three equal parts by trypsinization. One part was used to monitor the hSOAT protein expression by Western blot analysis, using the SOAT1 specific antibodies DM10 as the probe, as described ^12^, with the intensity of the WT hSOAT protein set as 1.0. The other two parts of the transfected cells were used to monitor the SOAT enzyme activity, by incubating the intact cells to 20 µl of ^3^H-oleate/fatty acid free BSA (7.5x 10^6^ dpm/µl) for 20 min. The amount of ^3^H-cholesteryl oleate produced was determined by the procedure previously described ^22,29^. Briefly, after ^3^H-oleate/BSA pulse, cells were rinsed with PBS, harvested in 1 mL per well of 0.1M NaOH, incubated at RT for 30 min. The cell homogenates were transferred to13x 100 mm size glass tubes; then neutralized by adding 67 µl of 3M HCl, and buffered with 50 µl of 1M K_2_HPO_4_. 80 µg per tube of non-radiolabeled cholesteryl oleate was added as carrier for identification purposes. Cellular lipids in each tube were extracted with 3 mL of CHCl_3_:MeOH at 2:1 followed by adding 1mL of H_2_O. The bottom chloroform phase that contained the lipid samples were dried under N_2_, then redissolved in 80 µl of ethyl acetate and spotted onto TLC plates (Anatech),with petroleum ether:ether:acetic acid (90:10:1) as the solvent system. After TLC, the plates were air-dried, the cholesteryl oleate bands (at Rf 0.9) were identified by briefly staining the TLC plates with iodine vapor. The cholesteryl oleate bands were scraped into scintillation vials and counted in a scintillation counter after addition of 3 mL per vial of Ecoscint O. The enzyme activity of each mutant hSOAT1 was estimated relative to that of WT hSOAT1, with the protein content of each mutant hSOAT1 normalized with that of the WT hSOAT.

### SOAT1 activity assay using NBD-cholesterol

The mixed micelles with 2.8 mM cholesterol/11.2 mM PC/18.6 mM taurocholate were prepared as described previously ^35^. The tetrameric or dimeric hSOAT1 enzyme was prepared in GDN detergent. First, 10 μl 2M KCl, 5μl 5% BSA, 1μl SOAT1 protein (A_280_=0.5), 5 μl micelles, 40 μl TBS buffer with 40 μM GDN were mixed with TBS with 0.5% CHAPS to reach the volume of 100 μl and incubated at 37°C for 2 min. In order to measure the IC_50_ of inhibitors, different concentrations of given inhibitors were added, as indicated. Then, 1.25 μl 0.2 mg/ml NBD-cholesterol (Sigma, N2161) solubilized in 35% β-cyclodextrin (Sigma, HZB1102) was added and the mixture was incubated at 37°C for 2 min. To start the enzymatic reaction, 1 μl 2.5 mM oleoyl-CoA (Sigma, O1012) was added and the reaction mixture was incubated at 37°C for 15 min. The reaction was terminated by adding 2:1 chloroform/methanol, the extract was separated on an HPLC column at 0.2 ml/min (Agilent, Poroshell HPH-C18, 2.7 μm) running in 100% ethanol and detected via fluorescence detector on an HPLC (SHIMADZU). NBD-cholesterol eluted at 1.3 min and its ester eluted at 1.9 min. The peak areas of the NBD-cholesteryl ester products and remaining NBD-cholesterol were integrated separately to obtain the relative ratio of NBD-cholesterol that was converted into NBD-cholesteryl esters.

### Nanodisc preparation

MSP2X was constructed by linking two MSP1E3D1 by PCR overlap extension, with Gly-Thr as the linker. The MSP2X gene was constructed into a pET vector, with N-terminal His_6_ tag and HRV 3C site. The MSP2X and MSP2N2 proteins were purified as described previously ^36^. The eluted SOAT1 protein in TBS buffer with 0.1% digitonin and 10 mM desthiobiotin from strep-tactin column was concentrated by a 100-kDa cut-off ultrafiltration device (Millipore) and exchanged into buffer without desthiobiotin. The SOAT1 protein was mixed with soybean polar lipids extract (SPLE, Avanti) and purified MSP (MSP2N2 for SOAT1 tetramer, MSP2X for SOAT1 dimer) at a molar ratio of SOAT1: MSP: SPLE = 1:7:100. For the hSOAT1 dimer in nanodisc that contained cholesterol, SPLE and cholesterol (at 4:1 ratio) were supplemented in GDN detergent micelles. The SOAT1 protein was mixed with micelles and purified MSP2X at a molar ratio of SOAT1: MSP2X: SPLE = 1:4:100. After incubating at 4°C for 30 min, Bio-beads SM2 (Bio-Rad) were added and rotated at 4°C for 1 h to initiate the reconstitution. Another batch of fresh bio-beads was added and rotated at 4°C overnight. The next day, the Bio-beads were removed and the mixture was loaded into a streptactin column to remove the empty nanodisc. The eluted hSOAT1 in the nanodisc was concentrated and cleaved by prescission protease to remove GFP tags. The nanodisc was centrifuged at 40,000 rpm for 30 min, and then loaded onto a superose-6 increase column running in TBS containing 0.5 mM TCEP. The collected fractions were detected by SDS−PAGE and peak fractions were concentrated to A_280_ = 1.2. Because the hSOAT1 tended to aggregate after nanodisc reconstitution, fractions of each peak were combined and concentrated for cryo-EM grids preparations and only fractions containing high ratio of SOAT1 dimer were used for data collection.

### Cryo-EM data collection

The nanodisc samples were loaded onto glow-discharged GiG R1/1 holey carbon gold grids (Lantuo) and plunged into liquid ethane by Vitrobot Mark IV (Thermo Fisher Scientific). Cryo-grids were screened by Talos Arctica electron microscope (Thermo Fisher Scientific) operated at the voltage of 200 kV using a Ceta 16M camera (Thermo Fisher Scientific). Optimal grids were transferred to Titan Krios electron microscope (Thermo Fisher Scientific) operated at the voltage of 300 kV, with an energy filter set to a slit width of 20 eV. Super-resolution movies (50 frames per movie) were collected with a dose rate of 5.4 e^−^/pixel/s using K2 Summit direct electron camera (Thermo Fisher Scientific) at a nominal magnification of 130,000 ×, equivalently to a calibrated super-resolution pixel size of 0.5225 Å, and with defocus ranging from −1.3 μm to −2.3 μm. All data acquisition was performed automatically using SerialEM ^37^.

### Cryo-EM image processing

For the CI-976 complex, super-resolution movie stacks were motion-corrected, dose-weighted and 2-fold binned by MotionCor2 1.1.0 using 9 × 9 patches ^38^. Micrographs with ice or ethane contamination were manually removed. Contrast transfer function (CTF) parameters were estimated using Gctf v1.06 ^39^. Particles were picked by Gautomatch (developed by Kai Zhang) and subjected to reference-free 2D classification. Unless otherwise stated, all classification and reconstruction were performed with Relion 2.0 ^40^. Initial model was generated by cryoSPARC ^41^ using the selected particles from 2D classification. The selected particles were further subjected to 3D classification using C1 symmetry. The particles selected from good 3D classes were re-centered and their local CTF parameters were determined using Gctf v1.06. These particles were further refined by cisTEM ^42^ using C2 symmetry imposed. The resolution estimation was based on the Part.FSC curve in cisTEM at FSC=0.143 cut-off. Local resolution estimation for hSOAT1 dimer and CI-976 complex was calculated using blocres ^43^. For the apo state, images were processed in the same way, except the finial refinement were done using Relion 3.0 ^44^ with a soft mask that excluded the MSP and lipids. The resolution estimations of apo state were based on the gold standard FSC of 0.143 cut-off after correction of the masking effect ^45^. Local resolution estimation for hSOAT1 dimer in apo state was calculated using Resmap ^46^.

### Model building and refinement

The sharpened map in the presence of CI-976 from cisTEM were converted into mtz file by Phenix ^47^. The model was manually built de novo in Coot ^48^. The assignment of transmembrane domain helices were based on their connectivity aided by the less-sharpened map. The register assignment and modeling building were based on the features of large aromatic side chains and partly aided by the further sharpened map. The manually built model was refined by Phenix ^47^. The sterol-like ligand in non-protein density A was built as cholesterol for visualization. The model in the presence of CI-976 were fitted into the map in the presence of BiSAS by Chimera ^49^ and further refined by Phenix.

### Quantification and statistical analysis

Global resolution estimations of cryo-EM density maps are based on the 0.143 Fourier Shell Correlation criterion ^50^. Fluorescence values were plotted versus the log of the concentration of inhibitor, and GraphPad Prism 6 was used to generate a curve fit with dose-response inhibition equation: Y=100/1+10^[Log(IC50-X) *HillSlope]^. IC_50_ values were calculated from the curve fit using Prism software. The number of biological replicates (N) and the relevant statistical parameters for each experiment (such as mean or standard error) are described in the figure legends. No statistical methods were used to pre-determine sample sizes.

---

**Fig. S1.**
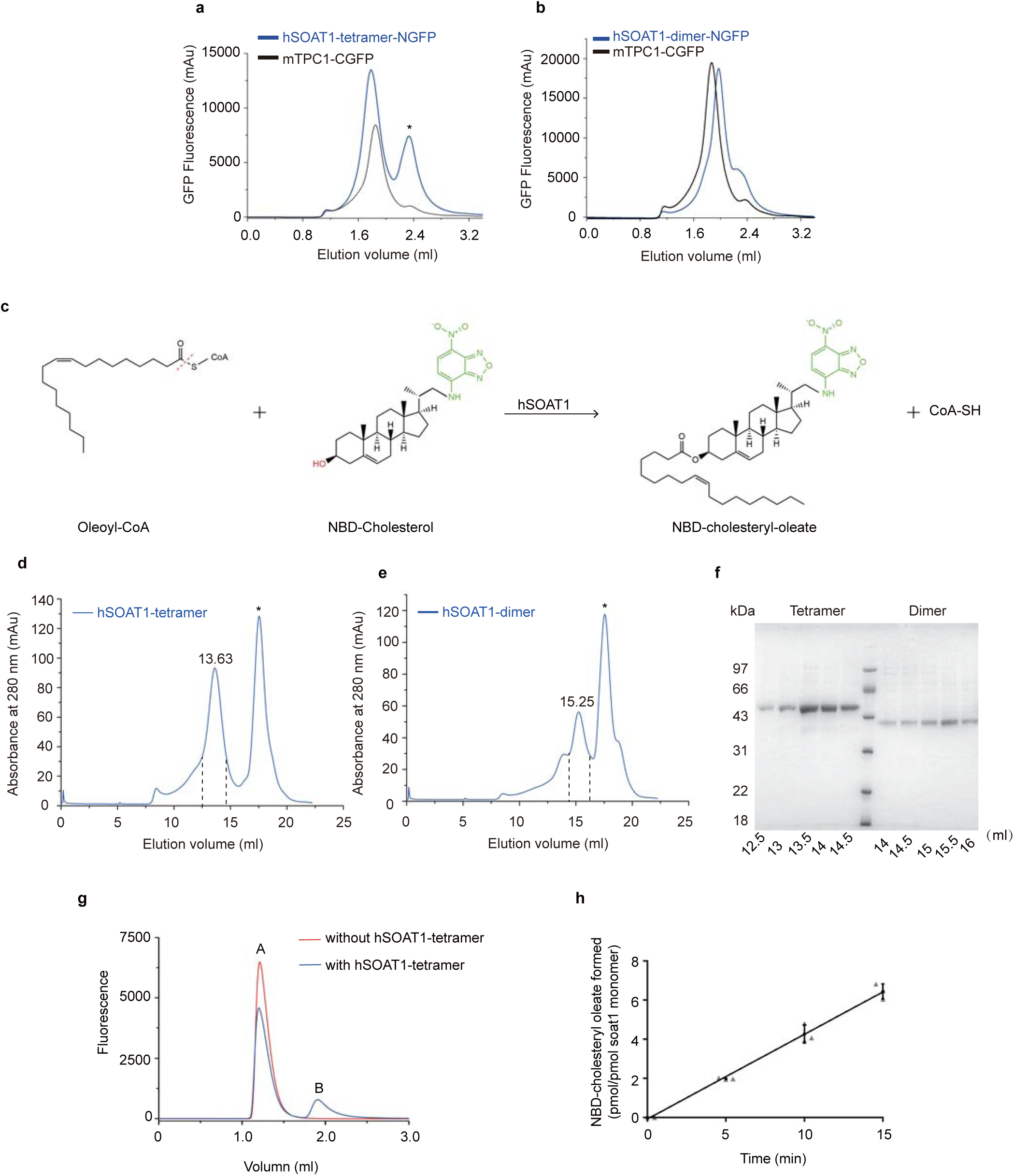
Characterization of human SOAT1 proteins. **a-b**, Fluorescence-detection size-exclusion chromatography (FSEC) traces of the N-terminal GFP tagged hSOAT1 tetramer and dimer on a Superose 6 increase column. The traces of C-terminal GFP tagged mouse TPCl were shown in black. The NGFP-hSOAT1 tetramer protein elutes at a position slightly earlier than the dimeric mTPC1-CGFP, while the NGFP-hSOAT1 dimer protein elutes later than the mTPC1-CGFP. An asterisk denotes the position of free GFP. **c**, The chemical reaction of hSOAT1 activity assay using NBD-cholesterol as substrate. The red dashed line indicates the bond that is broken during acyl-transfer reaction, the hydroxyl group that forms ester bond with the acyl group is highlighted in red. The NBD-fluorescent group is colored in green. **d-e,** The superose 6 elution profiles of hSOAT1 tetramer (**d**) and dimer (**e**), the fractions between the dashes were pooled and used for SDS-PAGE analysis. An asterisk denotes the position of GFP. **f**, The SDS-PAGE gel of purified hSOAT1 tetramer and dimer. **g**, The separation of NBD-cholesterol and NBD-cholesteryl-oleate by HPLC. Peak A is the free NBD-cholesterol. Peak B is the NBD-cholesteryl-oleate product. The fraction of NBD-cholesteryl-oleate product was calculated as area A/(area A+ area B). **h**, The reaction of hSOAT1 tetramer was linear with time within the first l5 min (Data are shown as means ± standard deviations, n = 3 biologically independent samples).

**Fig. S2.**
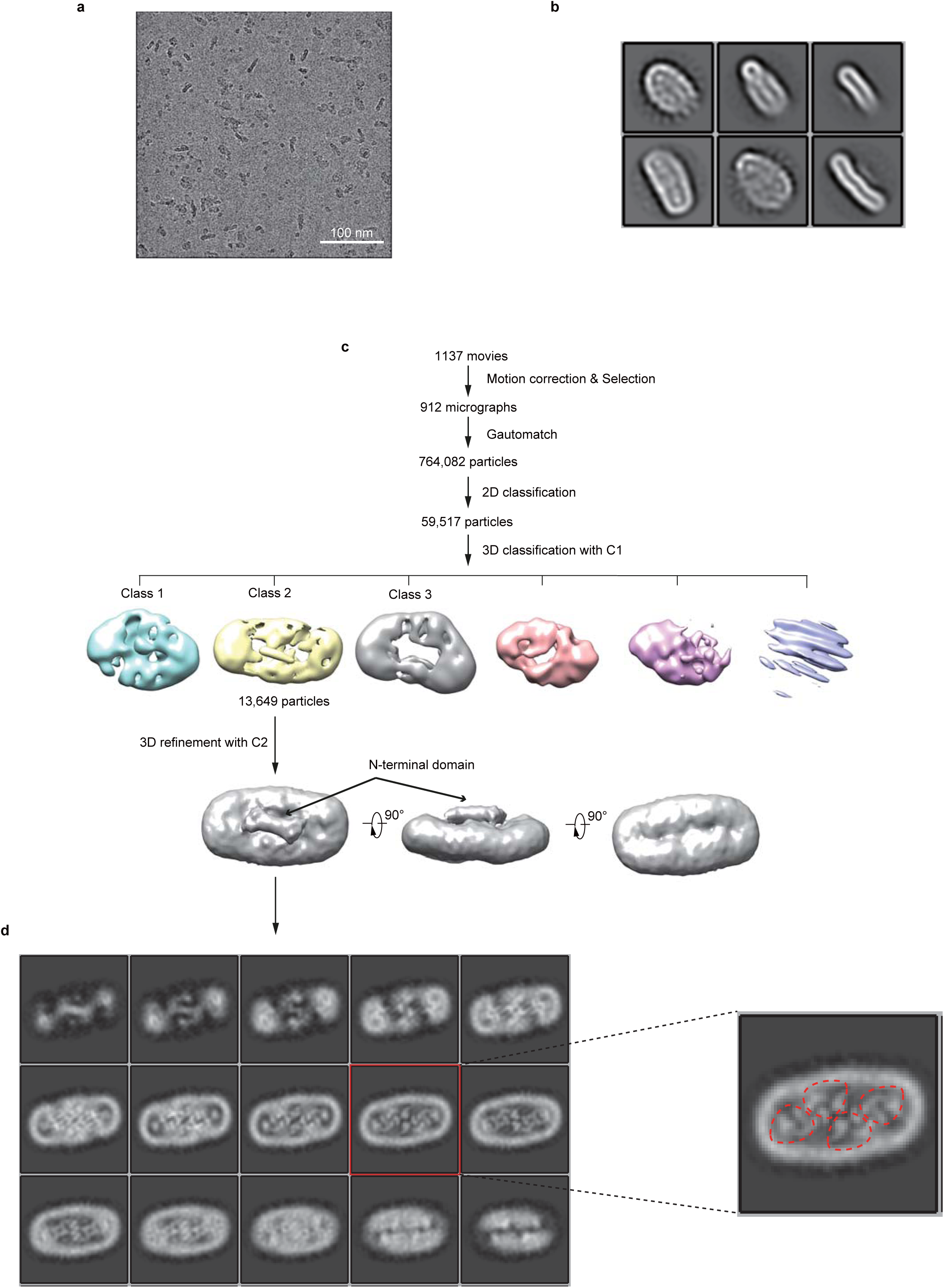
Cryo-EM image processing procedure of the hSOAT1 tetramer in digitonin detergent. **a**, Representative raw micrograph of hSOAT1 tetramer sample. **b**, Representative 2D class averages of the cryo-EM particles of hSOAT1 tetramer. **c**, Flowchart of the image processing procedure for hSOAT1 tetramer. **d**, The top-down slice view of the 3D density map after 3D refinement and postprocessing. The slice in red box is zoomed in for visualization. The red dashes circle each individual hSOAT1 monomer.

**Fig. S3.**
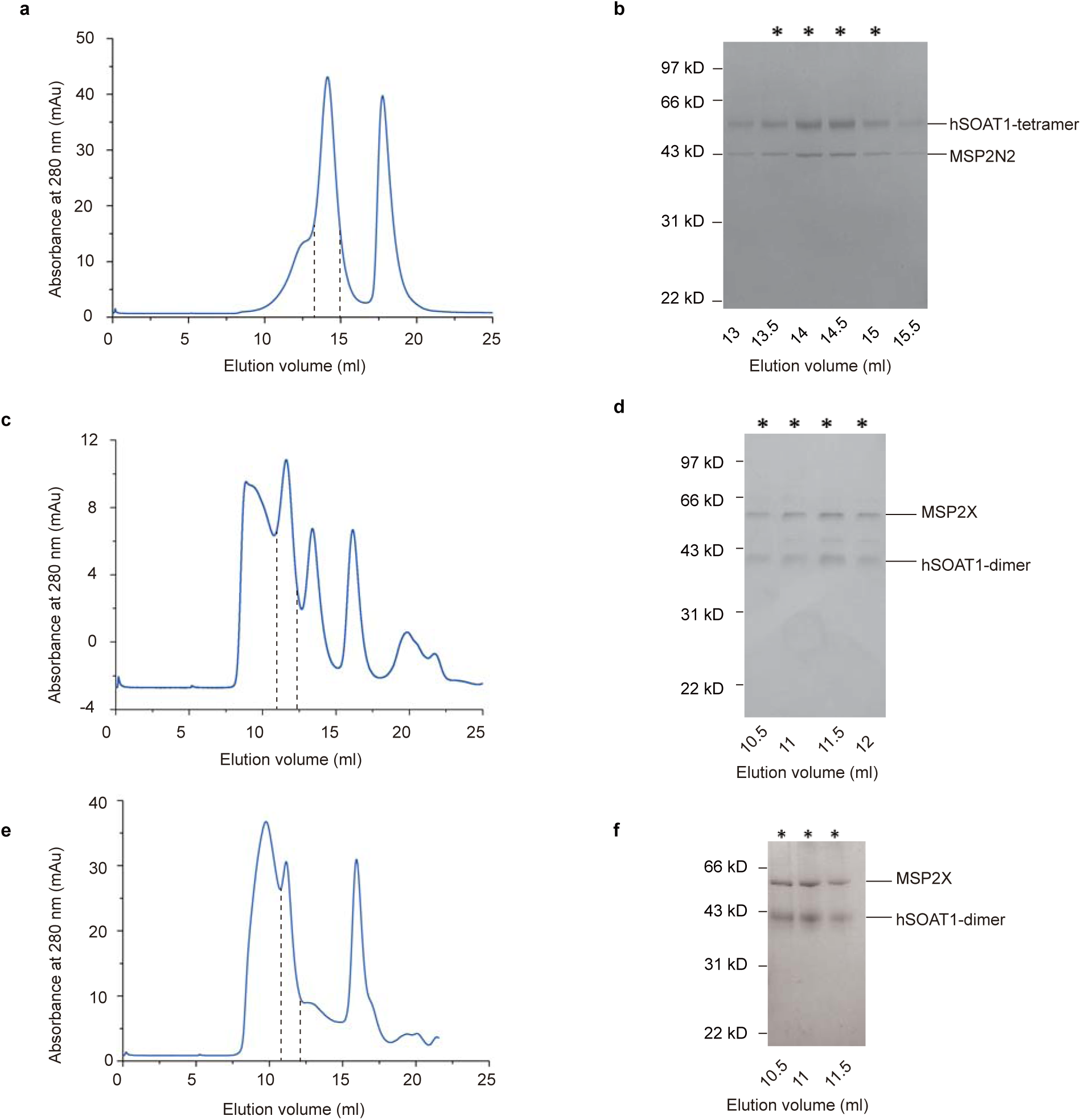
Purification of hSOAT1 tetramer and dimer nanodisc samples. **a**, Size exclusion chromatography (SEC) profile of the hSOAT1 tetramer nanodisc sample on Superose 6. The fractions between the dashes were pooled and used for cryo-EM analysis. **b**, hSOAT1 tetramer nanodisc samples of the indicated SEC fractions were subjected to SDS-PAGE and Coomassie blue staining. The asterisks denote the pooled fractions. **c-d**, Superdex 200 SEC profiles and SDS-PAGE of hSOAT1 dimer nanodisc in the presence of CI-976. **e-f**, SEC and SDS-PAGE results of hSOAT1 dimer nanodisc in the presence of cholesterol and BiSAS.

**Fig. S4.**
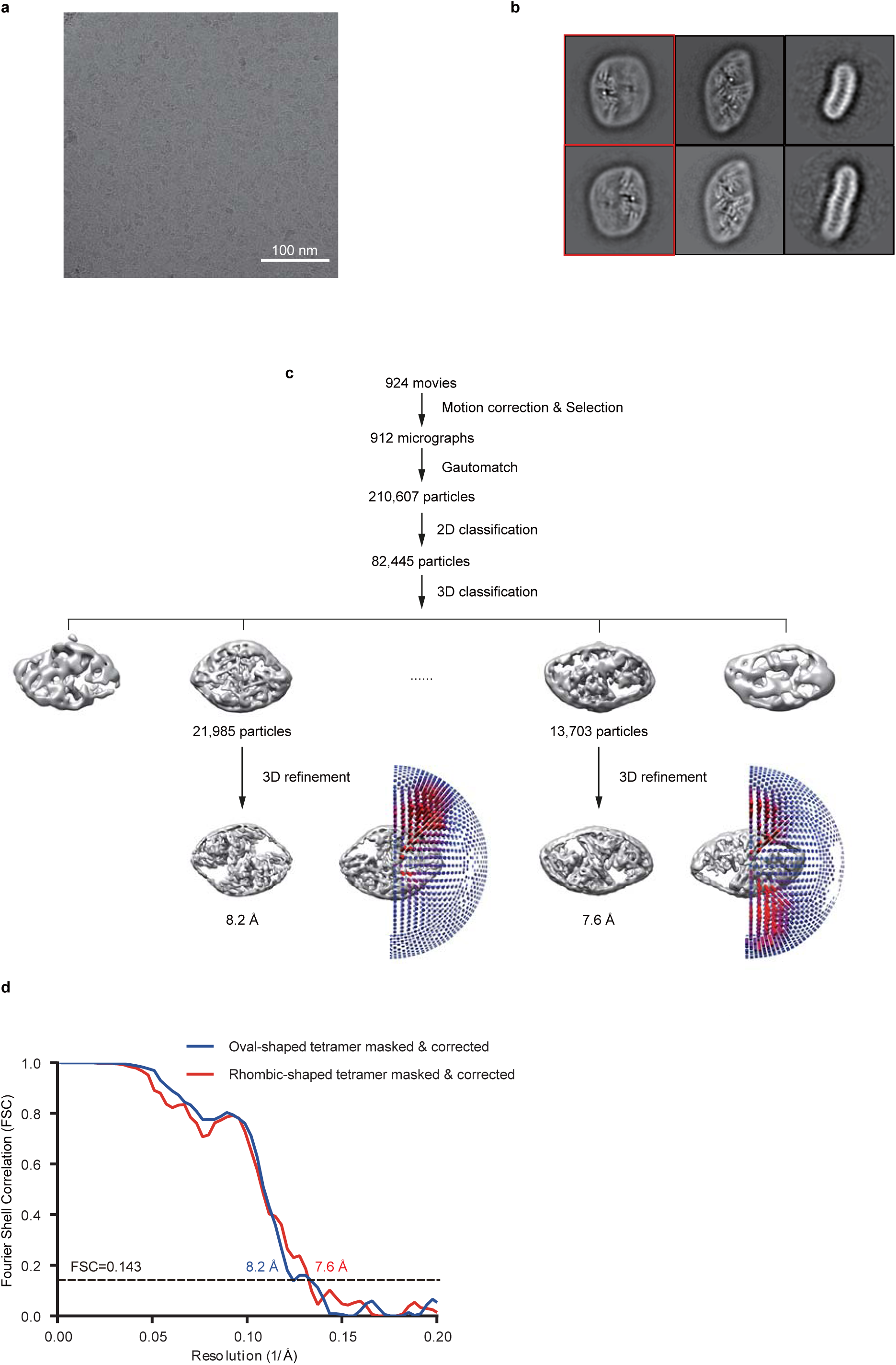
Cryo-EM image processing procedure of the hSOAT1 tetramer. **a**, Representative raw micrograph of hSOAT1 tetramer sample. **b**, Representative 2D class averages of the cryo-EM particles of hSOAT1 tetramer. The 2D class averages in red boxes show one clear dimer in adjacent to a blurry dimer, indicating the highly mobile interface between dimers. **c**, Flowchart of the image processing procedure for hSOAT1 tetramer. **d**, Gold-standard Fourier shell correlation (FSC) curves of the final refined maps for oval-shaped tetramer (blue line) and rhombic-shaped tetramer (red line). Resolution estimations (8.2 Å for the oval-shaped tetramer and 7.6 Å for the rhombic-shaped tetramer) are based on the criterion of an FSC cutoff of 0.143.

**Fig. S5.**
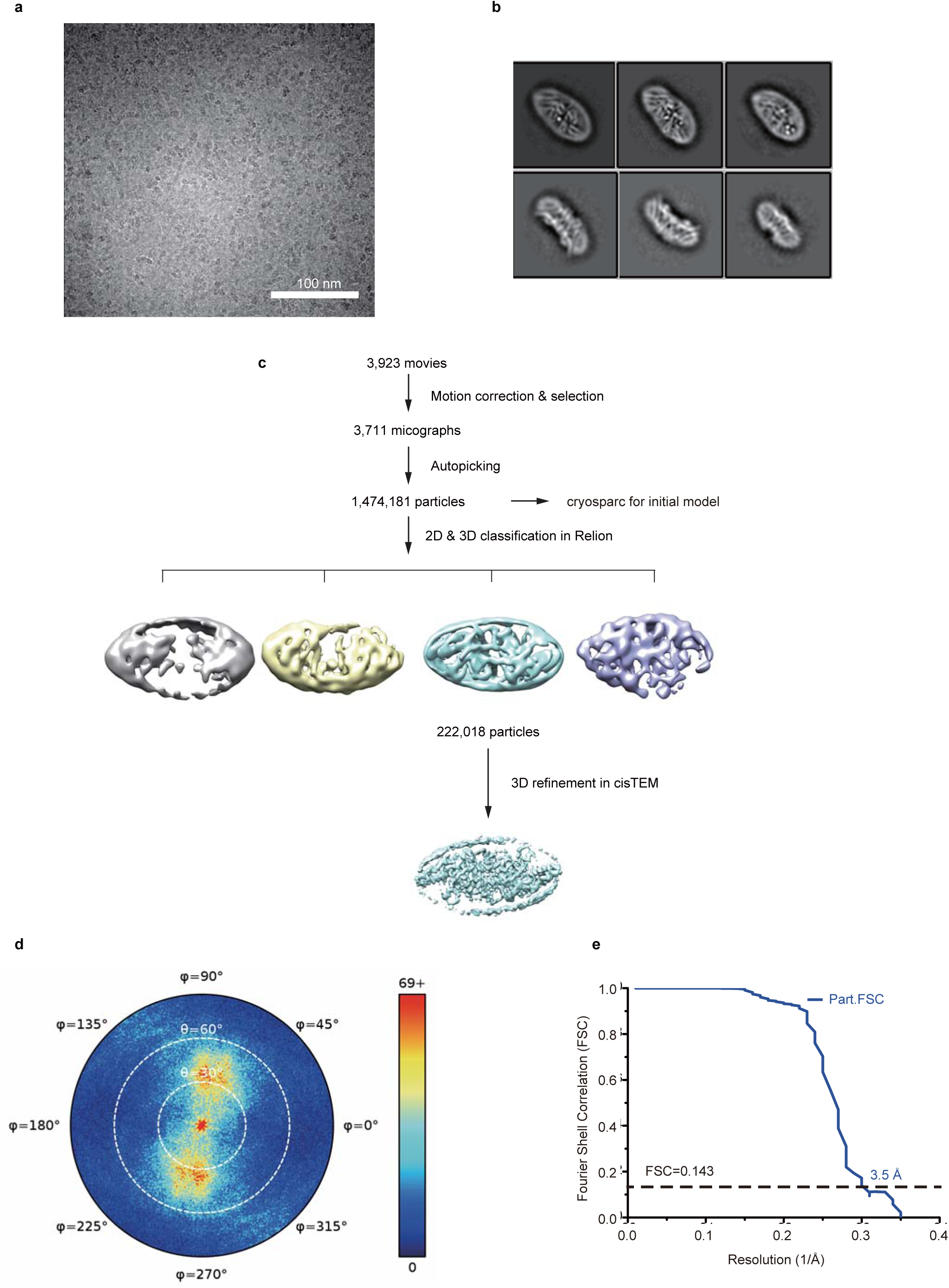
Cryo-EM image processing procedure of the hSOAT1 dimer in complex with Cl-976. **a**, Representative raw micrograph of hSOAT1 dimer. **b**, Representative 2D class averages of the cryo-EM particles of hSOAT1 dimer. **c**, Flowchart of the image processing procedure for hSOAT1 dimer. **d**, Angular distribution of the final reconstruction of hSOAT1 dimer. **e**, Gold-standard Fourier shell correlation (FSC) curve of the final refined map for hSOAT1 dimer. Resolution estimation (3.5 Å) is based on the criterion of the FSC cutoff at 0.143 in cisTEM.

**Fig. S6.**
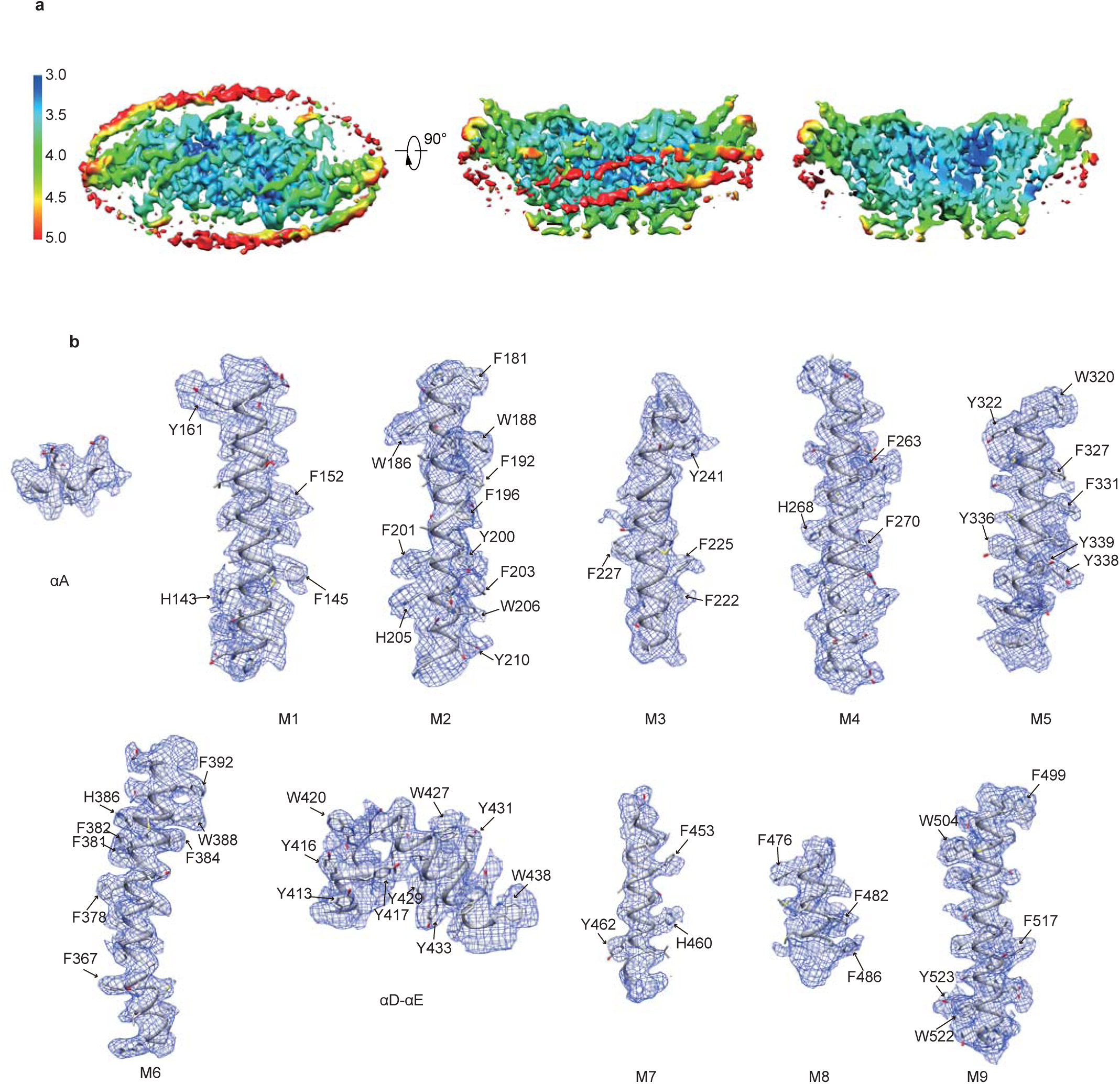
Electron density map of the hSOAT1 dimer in complex with Cl-976. **a**, Top view (left), side view (middle) and cut-away (right) representations of the hSOAT1 dimer cryo-EM density map colored according to the local resolution estimation. **b**, EM density segments (blue mesh) of the 9 transmembrane helices (Ml-M9), αA and αD-αE, .

**Fig. S7.**
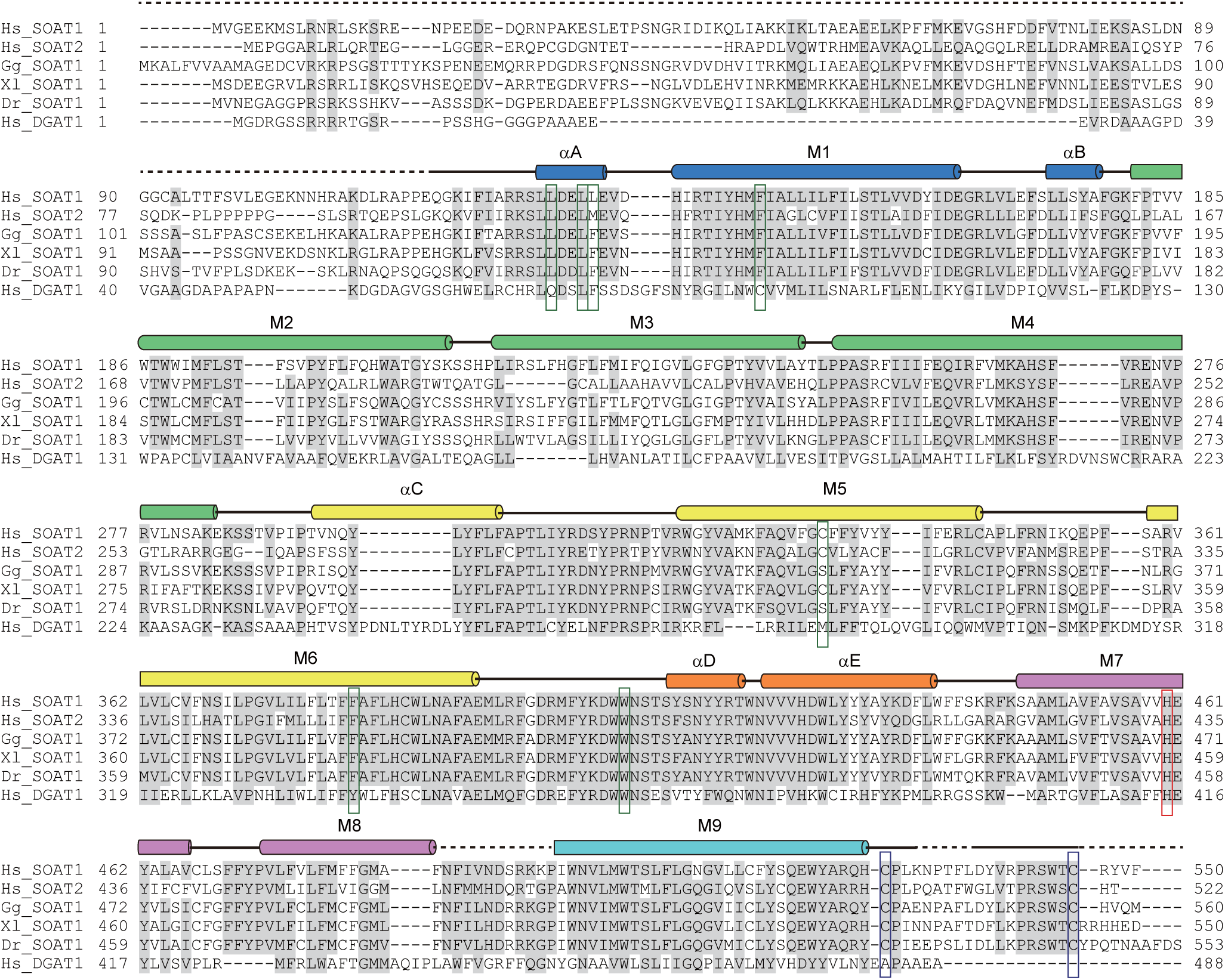
Sequence alignments of HsSOAT1, HsSOAT2, GgSOAT1, XlSOAT1, DrSOAT1 and HsDGAT1. The secondary structure elements are shown above the sequences (a-helices as cylinders, loops as lines and unmodeled residues as dashed lines). Conserved and highly conserved residues are highlighted in gray. Cylinders are colored in rainbow colors according to Fig. 3d. The active site H460 is boxed in red. Two cysteines forming the disulfide bond in the ER lumen are boxed in blue. Residues that interact with the putative sterol-like molecule are boxed in green. Hs: homo sapiens, Gg: Gallus gallus, Xl: Xenopus laevis, Dr: Danio rerio.

**Fig. S8.**
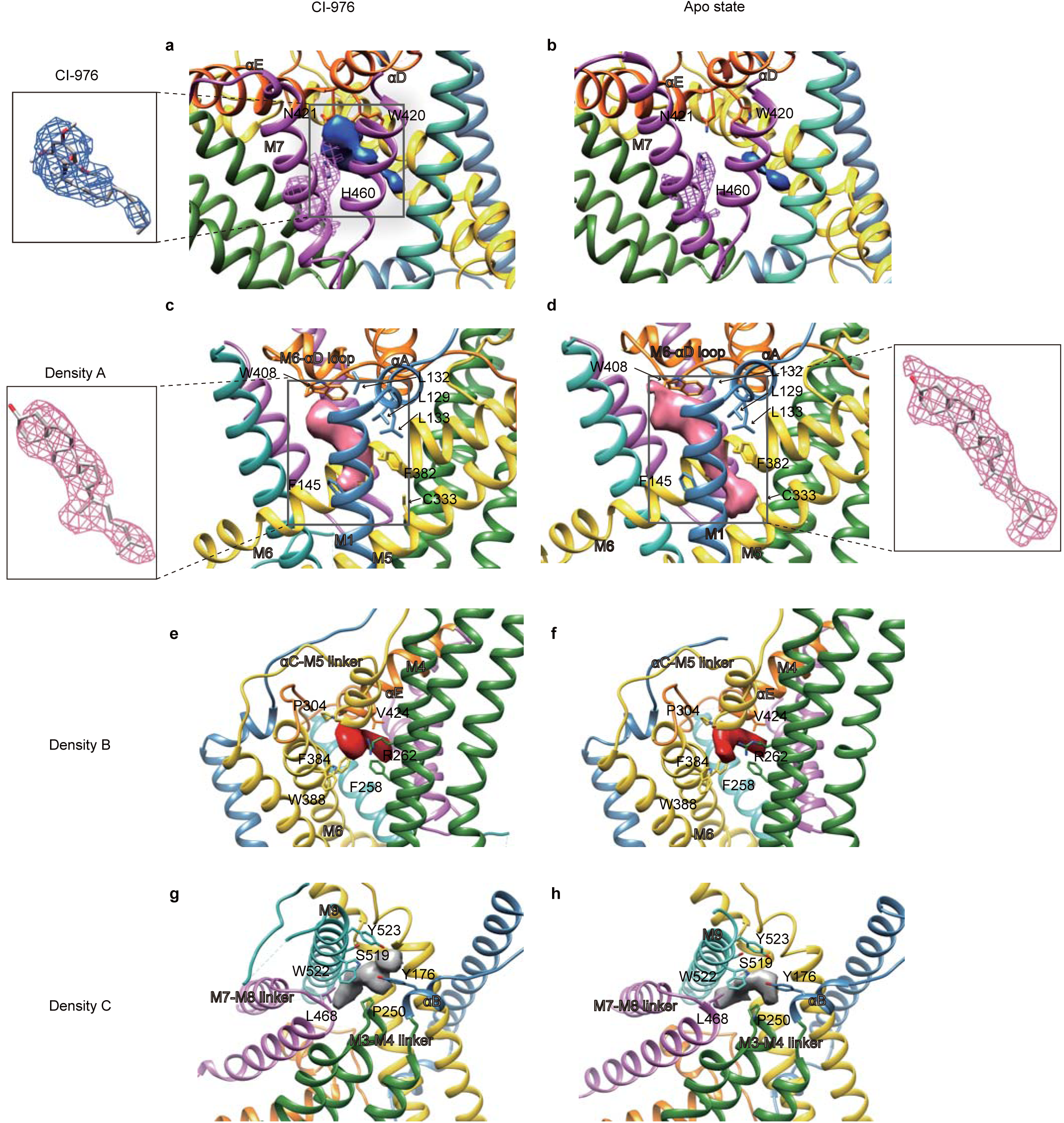
Electron density maps of bound ligands. **a-b**, Local EM densities inside the catalytic chamber in hSOAT1 dimer maps in complex with CI-976 (**a**) and in apo state (**b**). The inhibitor CI-976 density in (**a**) is shown as blue surface. The weak residual density in the BiSAS map is also shown as blue surface. The density of H460 side chains is shown in pink meshes at the same contour level as the ligand density in blue. **c-d**, The sterol-like densities (density A) in the maps of hSOAT1 dimer in complex with CI-976 (**c**) and in apo state (**d**) are shown in pink. The close-up view of the density with a sterol-like molecule inside is shown in boxes. **e-f**, The putative ligand densities (density B) in the maps of hSOAT1 dimer in complex with CI-976 (**e**) and in apo state (**f**) are shown in red. **g-h**, The putative ligand densities (density C) in the maps of hSOAT1 dimer in complex with CI-976 (**g**) and in apo state (**h**) are shown in grey.

**Fig. S9.**
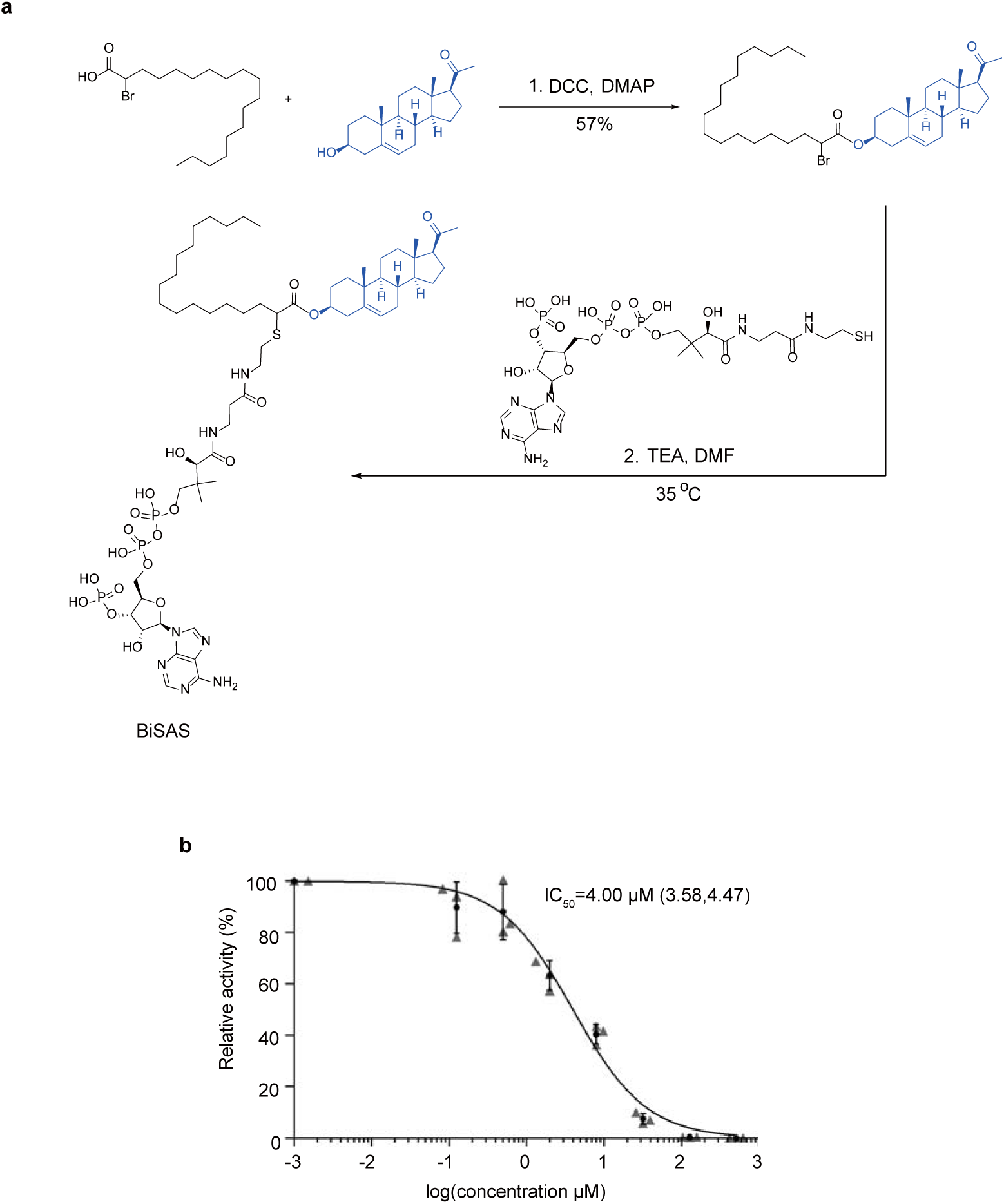
The chemical synthesis of BiSAS. **a**, Design and synthesis of BiSAS. **b**, Dose-dependent inhibition curve of hSOAT1 tetramer by BiSAS (The first data point is an artificial point. Data are shown as means ± standard deviations, n = 3 biologically independent samples).

**Fig. S10.**
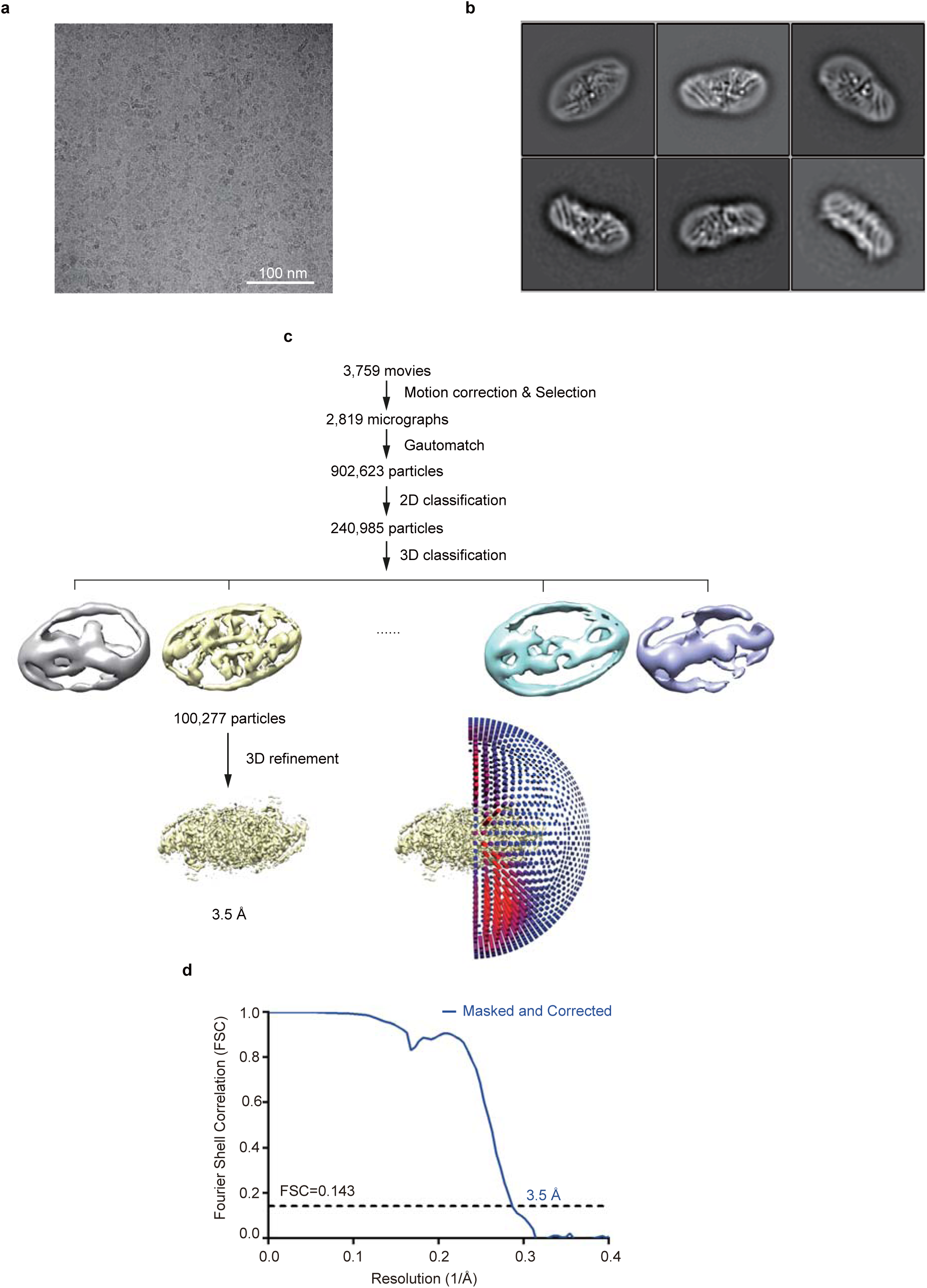
Cryo-EM image processing procedure of the hSOAT1 dimer in apo state. **a**, Representative raw micrograph of hSOAT1 dimer in apo state. **b**, Representative 2D class averages of the cryo-EM particles of hSOAT1 dimer in apo state. **c**, Flowchart of the image processing procedure for hSOAT1 dimer in apo state. **d**, Gold-standard Fourier shell correlation (FSC) curve of the final refined map for hSOAT1 dimer in apo state. Resolution estimation (3.5 Å) is based on the criterion of the gold-standard FSC cutoff at 0.143 in Relion.

**Fig. S11.**
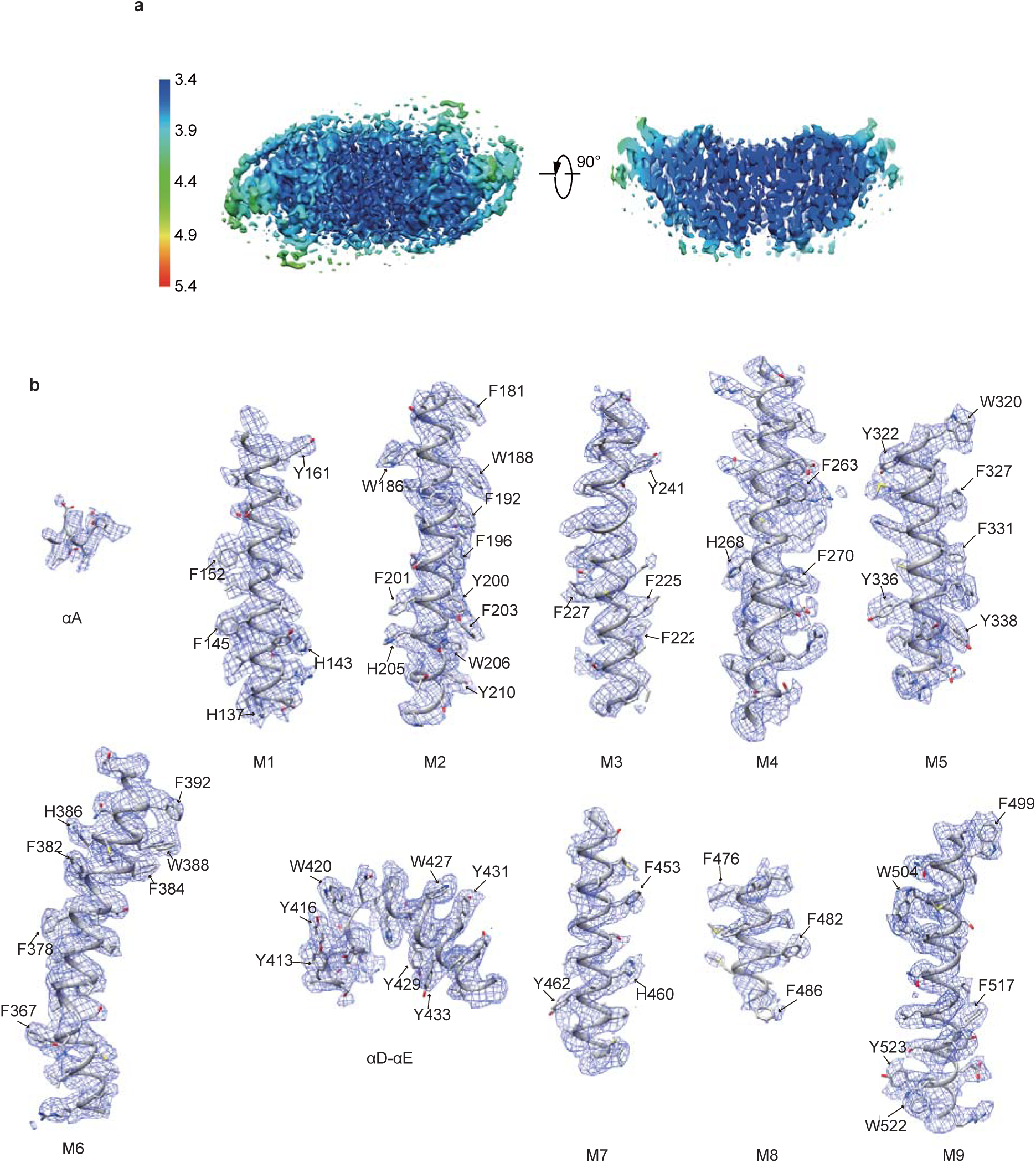
Electron density map of the hSOAT1 dimer in apo state. **a**, Top view and cut-away representations of the hSOAT1 dimer cryo-EM density map in apo state colored according to the local resolution estimation. **b**, EM density segments (blue mesh) of the 9 transmembrane helices (Ml-M9), αA and αD-αE, .

**Fig. S12.**
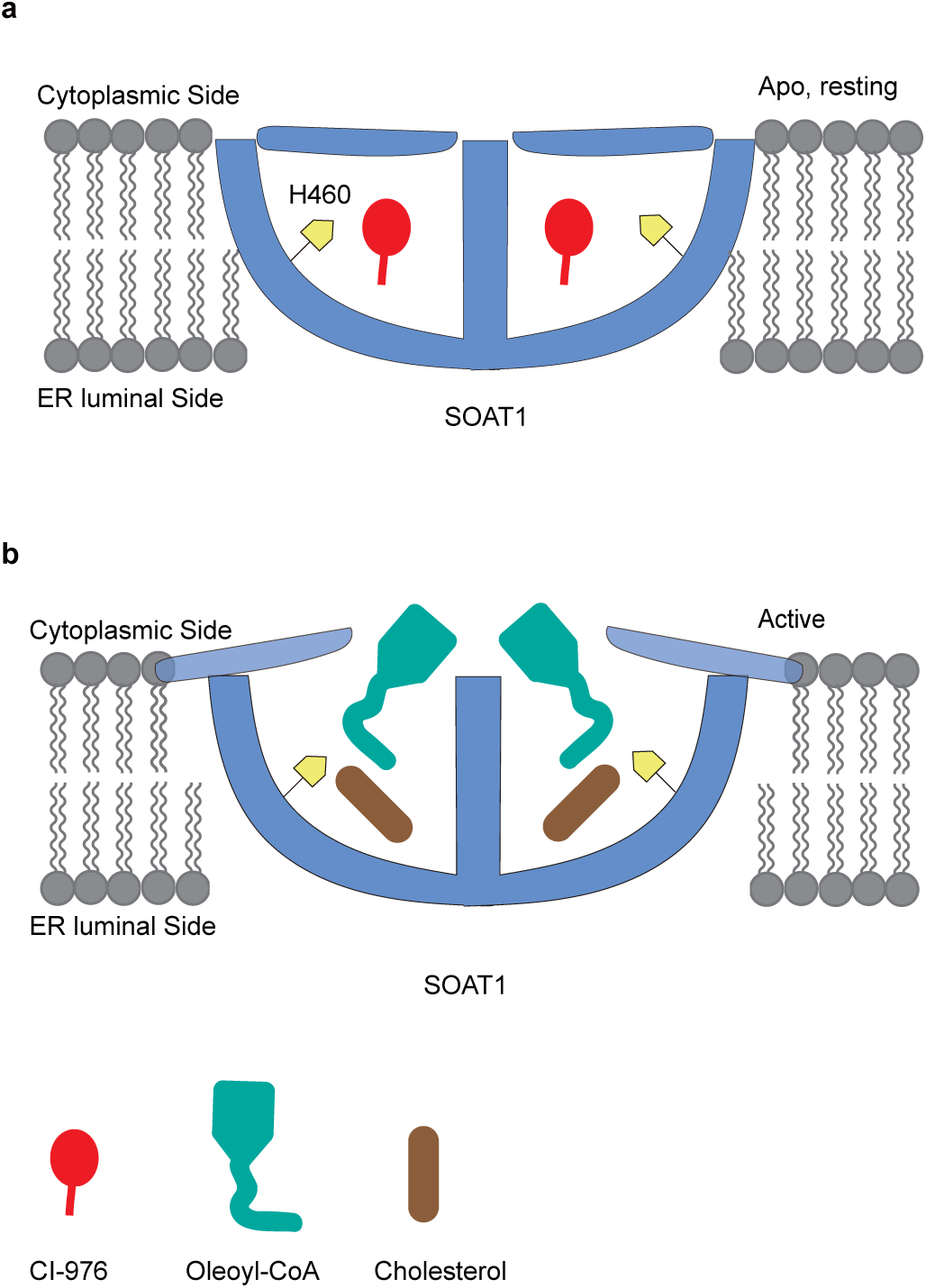
A working model to explain hSOAT1 activation. In the resting state, the putative catalytic residue H460 colored in yellow is less accessible to the acyl-CoA substrate. In the activation step, the lid of the reaction chamber is open; this step is required to activate the esterification reaction between acyl-CoA and cholesterol. For simplicity, only one SOAT1 dimer is shown.

